# Cell-intrinsic compliance mechanism enables release of tensile stress to prevent tissue rupture

**DOI:** 10.64898/2026.02.02.703250

**Authors:** Chun Wai Kwan, Shunta Sakaguchi, Michiko Takeda, Takefumi Kondo, Yu-Chiun Wang

**Affiliations:** Laboratory for Epithelial Morphogenesis, RIKEN Center for Biosystems Dynamics Research (BDR), Kobe, Japan; Graduate School of Biostudies, Kyoto university, Kyoto, Japan; Laboratory for Developmental Genome System, RIKEN BDR, Kobe, Japan; Graduate School of Medicine, Nagoya University, Nagoya, Japan

**Keywords:** Inter-tissue mechanical conflicts, Mechanical compliance, Extraembryonic tissue, Amnioserosa, Tensile stress, Junctional contractility, Tissue rupture, Optogenetics

## Abstract

Contractile forces are necessary to sculpt tissue structures and organ shapes during morphogenesis. In the early embryo, this presents an engineering challenge as the deforming tissue can be relatively large, while the forces thus generated might be poorly compartmentalized, generating tensile stresses that can cause damage if unmitigated. Here we show that during *Drosophila* gastrulation, the squamous morphogenesis of the extraembryonic amnioserosa functions mechanically to release tensile stresses yielded from the neighboring ectodermal convergent-extension and mesodermal invagination. Amnioserosa master regulator Zen transcriptionally silences *shroom* to endow it with low junctional actomyosin stresses, thereby ensuring high mechanical compliance. Loss of Zen, or targeted ectopic expression of Shroom in the amnioserosa using a novel optogenetic Gal4 system, leads to increased junctional myosin, thereby causing load-dependent tissue ruptures. Our data establish a previously unknown function for the *Drosophila* extraembryonic tissue, whereby cell-intrinsic mechanical compliance prevents tissue rupture to mitigate inter-tissue mechanical conflicts.

## Introduction

During animal development, embryos and organ primordia undergo a series of morphogenetic transformations to build precise and functional morphologies. Be it growth, deformation, rearrangement or migration, tissue morphogenesis requires active force generation. Given that morphogenetic processes do not occur in isolation, an increasing number of recent work in a variety of systems has highlighted the influence and contribution of geometry, boundary conditions, neighboring tissue dynamics, and external mechanical stresses^1–11^ on cell shape changes and tissue deformation. These data illustrate the importance of mechanical interplay when multiple morphogenetic processes take place concurrently.

The existence of inter-tissue mechanical interplay during morphogenesis implies that mechanical stresses generated in a tissue region may not be fully insulated or compartmentalized, or sufficiently dissipated within its spatial confine. As a result, undissipated morphogenetic stresses could potentially cause strains or lesions in the surrounding regions, which could be deleterious to development^10,11^. In the gastrulating embryo, multiple large-scale morphogenetic processes take place simultaneously within an interconnected epithelial monolayer that appears to behave as a mechanical continuum without apparent compartments or barriers that separate the regions of active morphogenesis. This raises the question of whether organisms might have evolved dedicated morphogenetic solutions that can mitigate the deleterious effects resulting from undissipated mechanical stresses.

One example of such a mechanical stress management program is cephalic furrow (CF) formation, which generates a transient epithelial fold during *Drosophila* gastrulation. The epithelial out-of-plane deformation associated with CF formation acts as a mechanical sink to release compressive stresses resulting from the collision of head and trunk tissues, preventing mechanical instabilities^10,11^. Whereas growth and tissue expansion produce pushing forces and can lead to the accumulation of compressive stresses, contractile behaviors can pull and yield tensile stresses in the neighboring tissues. If not properly released, such tensile stresses could cause mechanical lesions such as breakages, detachments or ruptures. Notably, two large-scale morphogenetic processes are driven by contractile forces during *Drosophila* gastrulation: 1) Ventral furrow (VF) formation is the process that internalizes the mesodermal precursor cells along the ventral midline, which leads to a net loss of ∼19 % of embryo surface area^12^. 2) Germband extension (GBE) is the convergent-extension event of the trunk ectoderm that causes the dorsal-ventral (D-V) extent of the tissue to narrow by ∼2 folds, thereby generating pulling force along the D-V circumference and and shearing stresses along the anterior-posterior (A-P) axis^6,13–15^. Biological tissues are able to release tensile stresses via either expansion or division^16,17^. A recent embryo-scale analysis reveals that cell division compensates for ∼65% of the lost surface area to all invagination events combined – including VF, CF, posterior midgut invagination (PMG) and dorsal fold (DF) formation^12^. It remains unclear, however, whether cell division plays the major role in releasing tensile stresses, or whether the embryo evolved other dedicated morphogenetic programs, such as tissue expansion, to mitigate deleterious effects that may arise due the accumulation of tensile stresses during *Drosophila* gastrulation.

In this study, we examined the developmental function of the morphogenetic process that generates the extraembryonic tissue of the *Drosophila* embryo, called amnioserosa (AS). We found that the specification of the AS fate endows the tissue with high mechanical compliance via a cell-intrinsic program that keeps the active actomyosin stresses low along the cell-cell junctions. As the processes of VF formation and GBE exert pulling forces on the AS, this cell-intrinsic program of low actomyosin stresses is essential to avoid the build-up of tensile stresses, thereby preventing the AS tissue from rupturing. The AS mechanical compliance stems from the transcriptional silencing of a scaffolding protein, Shroom, that promotes actomyosin contractility at the intercellular junctions. Our data suggest that during *Drosophila* gastrulation the AS squamous morphogenesis carries out a mechanical function that dissipates tensile stresses, thereby protecting the embryo from large-scale tissue rupture.

## Results

### AS apical elongation temporally follows surrounding morphogenetic events that generate contractile forces

The *Drosophila* extraembryonic tissue, the AS, is specified in a narrow strip of cells, spanning from ∼70% to ∼10% embryo length (EL; 0% EL refers to the posterior pole) along the A-P axis and corresponding to the dorsal-most 10% of cells at the dorsal midline^18,19^. As gastrulation commences, the AS tissue domain morphs from an elongated rectangular shape to a ‘saddle’ shape. This ‘saddle’ straddles the central portion of the dorsal midline, while extending down ventrally and bilaterally through the process of squamous morphogenesis, in which the individual cells undergo extreme apical-basal shortening from a columnar shape to a squamous shape, while the originally isotropic, hexagonal apical surfaces become anisotropic, taking on a highly elongated spindle shape^12,19,20^ (Fig. 1A-D, Movie S1).

**Figure 1.**
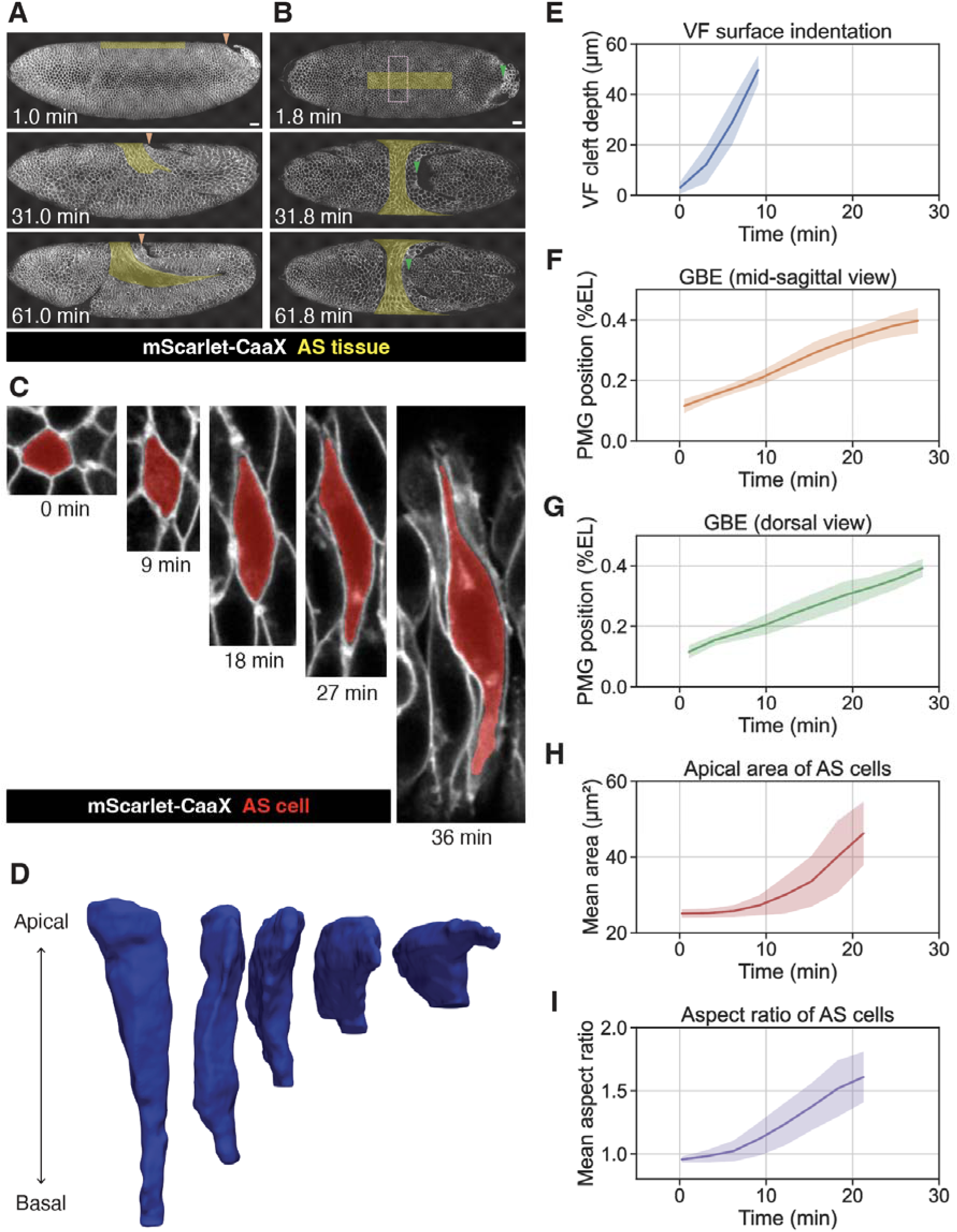
Morphodynamics of AS squamous morphogenesis in relation to VF formation and GBE. (A and B) Time-lapse images of an embryo expressing 3xmScarlet-CaaX showing the lateral (A) or the dorsal (B) surface of the embryo to illustrate the changes of AS tissue shape (highlighted with a yellow shade). Orange and green arrowhead, PMG position. Dotted rectangle in pale red, region of interest used for quantitation in (H and I). Time labels are elapsed time from the onset of VF formation. (C and D) A time series of the apical surface view of an AS cell (C, highlighted with a red shade) and its 3D rendered side view (D). Time labels are elapsed time from the onset of AS apical elongation. (E-I) Morphodynamics of VF cleft depth (E), PMG position in the mid-sagittal view from the lateral side (F), PMG position in the dorsal view (G), mean apical area of the AS cells (H) and mean aspect ratio of the AS cells (I). Time indicates elapsed time from the onset of VF formation. Scale bars, 20 µm. See also Movie S1.

To address the question of whether AS squamous morphogenesis plays a developmental role, we quantified its morphodynamics in relation to other morphogenetic events during gastrulation. *Drosophila* gastrulation begins with simultaneous onset of VF formation, PMG invagination and CF formation^11,14,21^. We temporally aligned all our live imaging datasets at the onset of VF formation and plotted metrics representing the progression of these morphogenetic events as a function of time. These include: 1) the increase of VF depth (Fig. 1E), 2) the anterior movement of the anterior edge of PMG for the progression of GBE (Fig. 1F, 1G), and 3) the apical surface elongation of the AS cells (Fig. 1H, 1I). The onset of AS elongation, as measured by the increase of apical surface area and aspect ratio, occurs ∼6 min after the onset of gastrulation. These data are consistent with a previous embryo-scale analysis^3^ and suggest that AS elongation temporally trails the initiation of VF invagination and GB convergent extension, while the ensuing VF deepening and GBE overlaps with the continuous increase of AS elongation. Thus, AS elongation could potentially be due to the external stresses resulting from VF and GBE morphogenesis.

### AS squamous morphogenesis requires both the Zen-dependent cell-intrinsic process and the external tensile stresses

The AS fate depends on the master regulator, *zerknüllt* (*zen*), which encodes a homeodomain transcription factor and is expressed in the presumptive AS tissue at the dorsal midline prior to the onset of gastrulation^22,23^. The dorsal midline expression of *zen* requires Decapentaplegic (Dpp) signaling, whose spatial patterning in turn depends on the graded activity of the Dorsal transcription factor along the D-V axis of the embryo^24–26^. Previous work suggests that cell shape change of AS depends on *zen*^20^. We thus asked whether AS elongation depends both on the Zen-dependent, cell-intrinsic program as well as on the external stresses generated by VF and GBE morphogenesis.

We first examined embryos lacking Zen activity (collectively abbreviated as the *zen^-^* embryos), using either a null mutation (*zen*^7^) or via knockdown of *zen* by the injection of double-stranded RNA targeting *zen* (*zen* RNAi). The dorsal midline cells undergo a reduced extent of apical surface elongation, consistent with previous work^20^, and the resultant saddle shape has a wider A-P span than in the wild-type (Fig. 2A, 2D, S1A, Movie S2). Quantitative analysis reveals that the dorsal midline cells in the *zen^-^* embryos do not undergo a full extent of elongation, suggesting that additional factors or forces, other than *zen*, are involved in promoting elongation (Fig. 2B, 2E). To test whether there might be additional genes downstream of the Dorsal gradient or Dpp signaling that act in parallel to *zen*, we examined embryos with a flattened Dorsal gradient (i.e. *cactus* RNAi) and *dpp* RNAi embryos. We found that these embryos showed similar phenotypes to the *zen^-^* embryos (Fig. S1A-C). Thus, these data do not support the presence of redundant genes. Instead, external stresses may contribute to AS elongation. In support of the possibility that external forces exert on the AS cells, we observed that the elongated cells in the ‘seat’ portion of the saddle show global ordering with their long axis aligned with each other and to become parallel to the embryonic D-V circumference^3^ (Fig. 2C) and that such global ordering persists in the *zen* RNAi embryos (Fig. 2F). These data suggest the existence of Zen-independent, D-V oriented external stresses that act on the AS cells.

**Figure 2.**
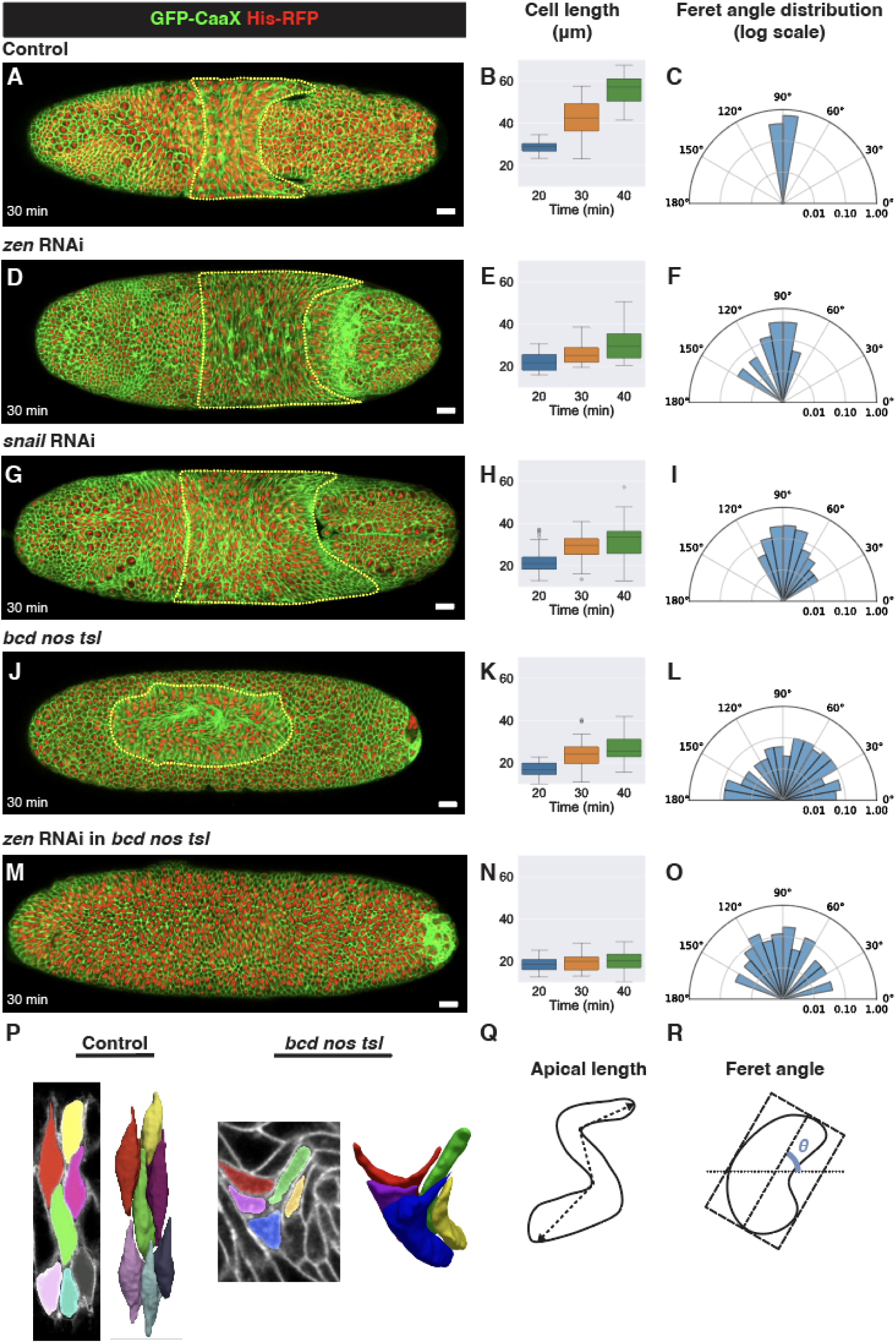
AS apical elongation and global alignment are coupled to external stresses. (A, D, G, J and M) Dorsal view of a control (A, n=5 embryos), *zen* RNAi (D, n=3 embryos), *snail* RNAi (G, n=3 embryos), *bnt* mutant (J, n=4 embryos) or *bnt* mutant with *zen* RNAi (M, n=3 embryos) embryo expressing GFP-CaaX and His-RFP, 30 min after the onset of AS apical elongation. Yellow dotted outline, the AS or the corresponding dorsal region that contains cells that are apically elongated. Scale bars, 20 µm. (B, C, E, F, H, I, K, L, N and O) The apical cell length (geodesic diameter, see also Q) over time and the percentage distribution of the Feret angles (see also R) of the AS cells at 30 min in control (B and C, n=51 cells), *zen* RNAi (E and F, n=84 cells), *snail* RNAi (H and I, n=146 cells), *bnt* mutants (K and L, n=150 cells) or *zen* RNAi in *bnt* mutants (N and O, n=68 cells). (P) Segmentation (left) and 3D rendered apical view (right) of representative AS cells in a control or *bnt* mutant embryo. (Q and R) Schematics representations for how the apical cell length (geodesic diameter) and the Feret angle were measured. See also Figures S1, S2, Movies S2 and S3.

Previous work suggests that VF formation and GBE drive a ventral-ward tissue flow^3,11,27,28^. Given that the onset of VF formation and GBE precedes AS elongation (Fig. 1E-I), they thus could exert pulling forces on the AS cells. To test this, we disrupted VF formation using *snail* RNAi and separately used *bicoid nanos torso-like* (*bcd nos tsl* in short as *bnt*) triple mutant embryos to block GBE, as spatial patterning along the entire A-P axis and PMG are completely lost in these embryos, and GBE is fully abolished^15,29^. Both *snail* RNAi and *bnt* mutants show reduced AS elongation, as compared to the wild-type (Fig. 2G, 2H, 2J, 2K, S2A, Movie S2, S3), suggesting that both VF formation and GBE generates pulling forces to promote AS elongation. Strikingly, the elongated cells do not align with the D-V circumference in *bnt* mutants, but display randomized orientations, and the AS tissue fails to adopt a saddle shape, but retain an elongated oval shape, contrasting with *zen* or *snail* RNAi (Fig. 2F, 2I, 2L). These data are consistent with a recent report that analyzes the ordering of the AS cells^30^ and suggest that shearing stresses generated during GBE promotes global alignment of the AS cells.

Finally, we eliminated both the intrinsic and extrinsic forces that drive AS elongation. Specifically, we performed *zen* RNAi knockdown or *zen* and *snail* double RNAi knockdown in *bnt* mutants. We found that the dorsal midline cells fail to elongate at all, and consequently it is not possible to identify a region of apical elongation and cell alignment (Fig. 2M-O, S2A-C, Movie S2, S3), confirming that both the Zen-dependent cell-intrinsic program and the externally applied forces contribute to AS elongation. Thus, full-scale elongation of the AS cells require both the external pulling forces (VF + GBE) and the Zen-dependent cell-intrinsic program, while GBE additionally establishes boundary conditions and flow shear stresses that mold the AS into a saddle shape, while driving the global ordering of the spindle-shaped AS cells.

### The AS fate enables the release of tensile stresses to prevent tissue rupture

That AS elongation requires external pulling forces suggests that the AS cells are under external mechanical load, raising the possibility that AS squamous morphogenesis functions to release tensile stresses generated by VF formation and GBE. This hypothesis predicts that blockage of the cell-intrinsic program of elongation could lead to abnormalities linked to unreleased tensile stresses. To test this, we imaged long time-scale morphogenetic dynamics of the dorsal midline tissue in the *zen^-^* embryos and found that 32∼56 minutes after the onset of cell elongation, tissue tears emerge at the dorsal midline, culminating in large-scale ruptures (Fig. 3A-E, Movie S4, Data S1). The initial tears can be seen occurring at stochastic locations in the dorsal midline tissue. These tears occur as a result of the separation of cell-cell junctions. They are additionally associated with bursts of membrane extensions that may be related to attempts to close the wound. While some of these initial tears apparently healed, other tears ultimately rupture the tissue, forming large-scale holes. We observed complete penetrance of the rupture phenotype in the *zen^-^* embryos, and similar phenotypes in *cactus* and *dpp* RNAi embryos (Fig. S1D-F, Data S1). These data reveal that the Zen-dependent cell-intrinsic program prevents tissue rupture.

**Figure 3.**
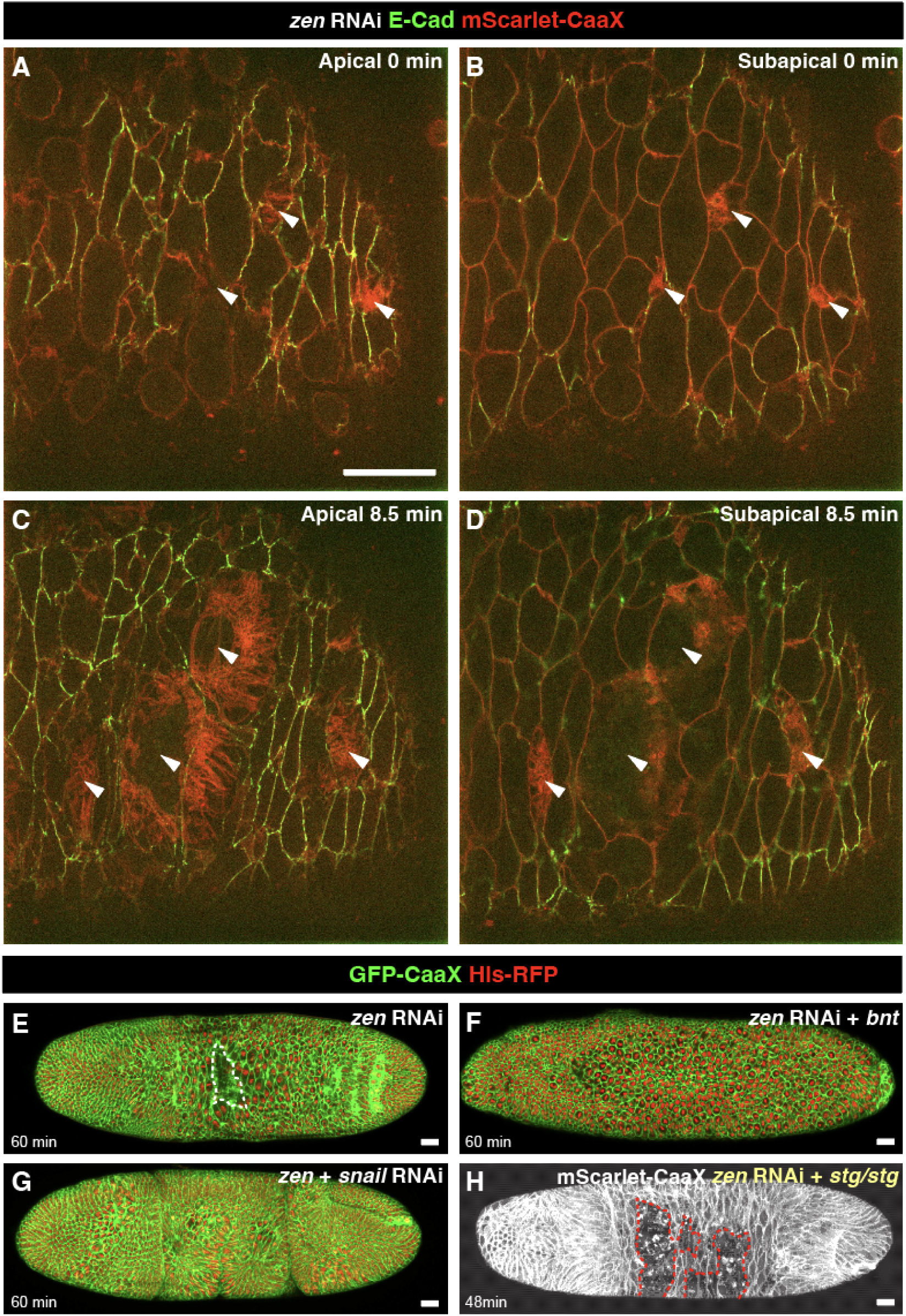
*zen* RNAi leads to tissue rupture in the presence of external stresses. (A-D) Dorsal view of a *zen* RNAi embryo expressing E-Cad-3xGFP and 3xmScarlet-CaaX (n=4) showing the onset (A and B) or 8.5 min after the onset (C and D) of tissue rupture in a single z-slice view at an apical (A and C) or subapical (B and D) plane. White arrows, sites of tissue rupture. (E-G) Dorsal view of a *zen* RNAi embryo expressing GFP-CaaX and His-RFP showing tissue rupture (E, n=3), or *zen* RNAi embryos with reduced external stresses showing the suppression of tissue rupture (F, *bnt* mutant, n=4; G, *snail* RNAi, n=3). White dotted outline, area of tissue rupture. (H) Dorsal view of a *zen* RNAi embryo also mutant for *stg* (n=2) showing tissue ruptures are not suppressed by the loss of cell divisions. The membrane marker is 3xmScarlet-CaaX. Red dotted outline, area of tissue rupture. Time labels are elapsed time from the onset of AS apical elongation. Scale bars, 20 µm. See also Figure S1, Movies S2 and S4.

To test whether tissue rupture in the *zen^-^* embryos is linked to tensile stresses generated during VF formation and GBE, we blocked these two processes using *snail* RNAi and *bnt* mutants, respectively. We found that indeed the ruptures are suppressed (2/3 *snail* RNAi embryos and 4/4 *bnt* mutant embryos; Fig. 3E-G, Movie S2). Thus, pulling forces exerted by VF formation and GBE leads to the accumulation of tensile stresses that rupture the dorsal tissue if the Zen-dependent cell-intrinsic program is absent.

We next sought to gain mechanistic insights into how Zen programs the AS cells to prevent tissue rupture. Previous work has shown that the AS fate is associated with the exit of the cell cycle and suggested that Zen-dependent transcriptional repression of the mitotic driver *cdc25/string* underlies it^31–33^. We confirmed and observed in the *zen^-^* embryos the resumption of mitosis in the normally mitotically quiescent AS (Movie S2). We thus wondered whether mitosis weakens cell-cell adhesion, rendering the tissue less resistant to external stresses and thus prone to rupture. To test this, we removed the activity of *cdc25/string* in *zen* RNAi embryos to block mitosis. We found, however, that the embryos still undergo tissue ruptures (Fig. 3H, Movie S2). These data suggest that tissue ruptures in the *zen^-^* embryo do not result from mitosis-driven weakening of cell adhesion.

### Mechanical compliance of the AS cells is linked to low junctional myosin

We next asked whether Zen-dependent mechanical properties of the AS cells account for the ability of the tissue to withstand tensile stresses. We examined two key determinants of epithelial mechanics: cell-cell adhesion and actomyosin contractility^34,35^. Quantitative analysis of the level of E-Cadherin (E-Cad), a core component of adherens junctions, reveals decreased intensity in the *zen^-^* embryos during dorsal cell elongation, as compared to the wild-type (Fig. 4A, 4B, 4A’, 4B’, 4C, 4D). In contrast, junctional myosin levels are high in the *zen^-^* embryos, contrasting with the wild-type embryos, which show low levels of junctional myosin along cell-cell junctions between the AS cells, except for the vertices^20^ (Fig. 4A’’, 4B’’, 4E, 4F, Movie S5). These data suggest that tissue rupture in the *zen^-^* embryo could be attributed to either low intercellular adhesion, or high junctional actomyosin contractility, or both.

**Figure 4.**
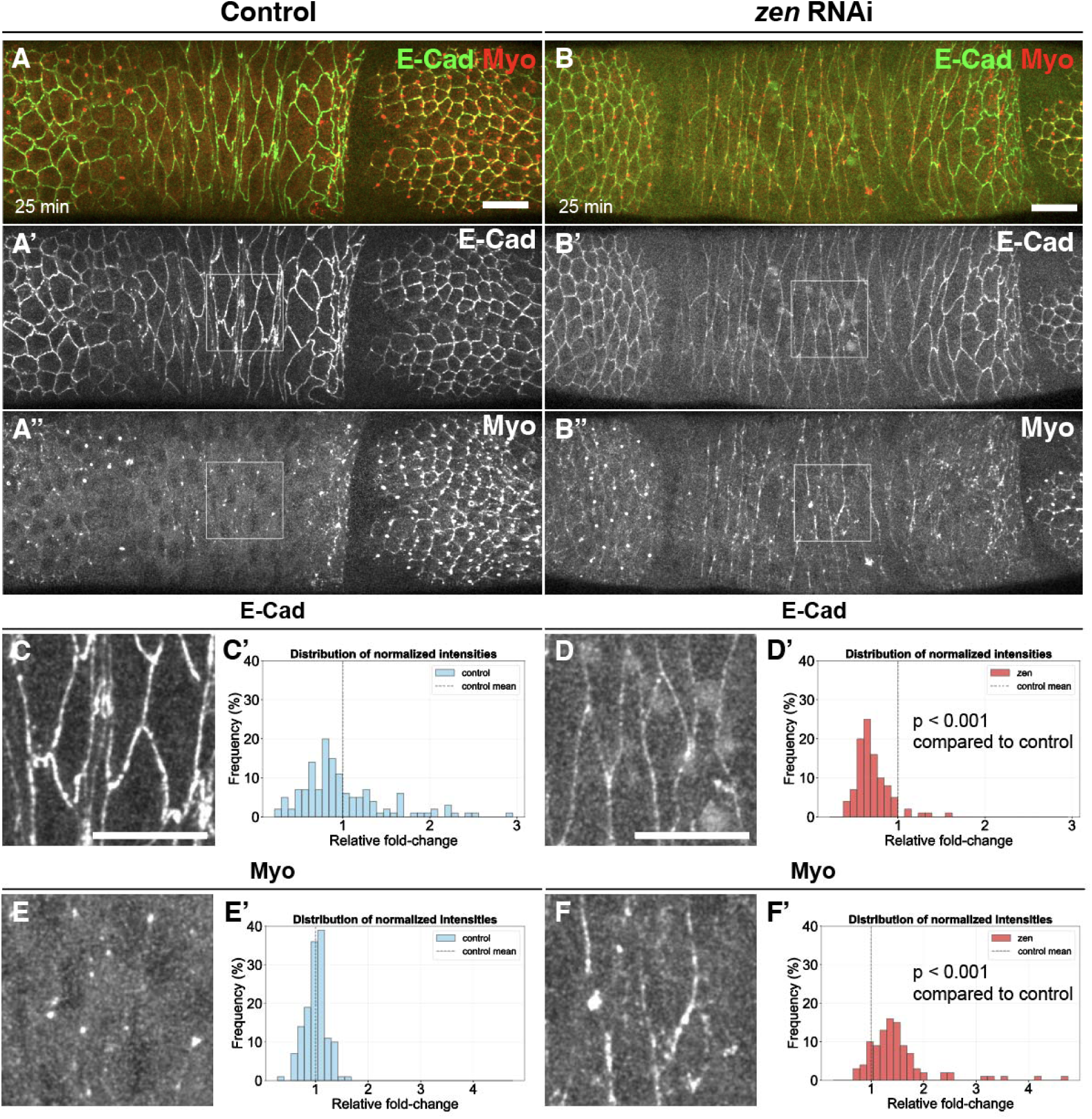
Zen lowers junctional myosin in the AS cells. (A and B) Dorsal view of a control (A, n=3) or a *zen* RNAi (B, n=3) embryo expressing E-Cad-3xGFP (Green in A and B; A’ and B’) and MyoII-3xmKate2 (Red in A and B; A’’ and B’’), 25 min after the onset of AS apical elongation. White boxes outline a representative region of the AS enlarged in C-F. (C-F) Enlarged view of regions outlined in white in A’, A’’, B’ and B’’, highlighting a decrease in E-Cad (C and D) and an up-regulation of junctional myosin (E and F) in *zen* RNAi, as compared to control. Scale bars, 20 µm. (C’-F’) Histogram showing the distribution of E-Cad and myosin junctional intensities, 25 min after the onset of AS apical elongation, quantified from (C-F) in control (C’ and E’, n=117 junctions) or *zen* RNAi (D’ and F’, n=83, junctions). Mann-Whitney U test; p < 0.001. See also Figure S2 and Movie S5.

We noticed that when the tissue tear first appears in the dorsal tissue in the *zen^-^* embryo, the dorsal cells are already stretched, while there is a lack of overt tissue flow away from the dorsal midline as tears enlarge to form large-scale holes (Movie S4). These observations suggest a potential similarity between our *in vivo* system and a recently reported *in vitro* epithelial model for tissue rupturing. In particular, the rupture we observed resembles a regime shown in the *in vitro* system, wherein a pre-stretched tissue with a constant strain forms internal tears when tissue-intrinsic tension increases and eventually reaches the tensile strength of the tissue^36^. The similarity suggests the possibility that high junctional actomyosin contractility is the dominant mechanical parameter in the *zen^-^* embryo. On the one hand, high junctional contractility during the initial loading stage of external tensile stresses likely limits the deformability of the dorsal cells (Fig. 2D, 2E); on the other hand, rising junctional contractility may ultimately cause the tissue to rupture when it exceeds the tensile strength of the tissue, even after VF invagination has completed and the dorsal cells are not stretched further.

We first tested whether AS cell deformability may be limited by increased contractility in the *zen^-^* embryo. Myosin contractility requires phosphorylation of its regulatory light chain (MRLC), while dephosphorylation of MRLC by protein phosphatase 1 inactivates it^37,38^. To increase myosin contractility, we used Calyculin A, a chemical inhibitor of serine/threonine protein phosphatases^39^. Injection of Calyculin A into *bnt* mutant embryos results in a complete loss of AS elongation, resembling the effect of *zen* RNAi on the *bnt* embryo (Fig. S2D-F). These results establish a causal link between myosin contractility and AS cell deformability, and suggest that increased myosin contractility in the *zen^-^* embryos confers low compliance to the AS cells when under a mechanical load.

### Single-cell RNA seq identifies Shroom as a transcriptional target silenced by Zen and required for low junctional myosin in the AS

The transcriptional targets of Zen that control the cellular and mechanical behaviors of the AS cells have not been identified previously. To elucidate the Zen-dependent intrinsic mechanism that confers mechanical compliance, we re-analyzed previously published single cell RNA-seq (scRNA-seq) datasets to identify candidate target genes downstream of Zen^40^. Through comparing the transcriptomes of the AS cells in the trunk region with those of the dorsal ectoderm, we identified 65 genes that are upregulated (AS-high, Data S2) and 33 genes downregulated (AS-low, Data S3) in the AS (Fig. 5A). As expected, *zen* was among the AS-high genes, so are several dorsally expressed genes known to require Zen (Fig. 5B). In search of putative Zen targets that could account for the low levels of junctional myosin in the AS cells, we considered *shroom* (*shrm*), which is downregulated in the AS, as compared to the ectoderm (Fig. 5B-D). Shrm is a scaffolding protein that contains binding sites for actin and Rho-kinase, and known to be enriched along adherens junctions by virtue of its actin binding capability^41–43^. Shrm recruits Rho-kinase to activate MRLC, thereby elevating contractility at adherens junctions. We proposed that Zen transcriptionally represses *shrm* in the AS cells to maintain low levels of junctional myosin contractility and cell-intrinsic stresses, thereby allowing the AS cells to be compliant when the tissue is under high external tensile stresses.

**Figure 5.**
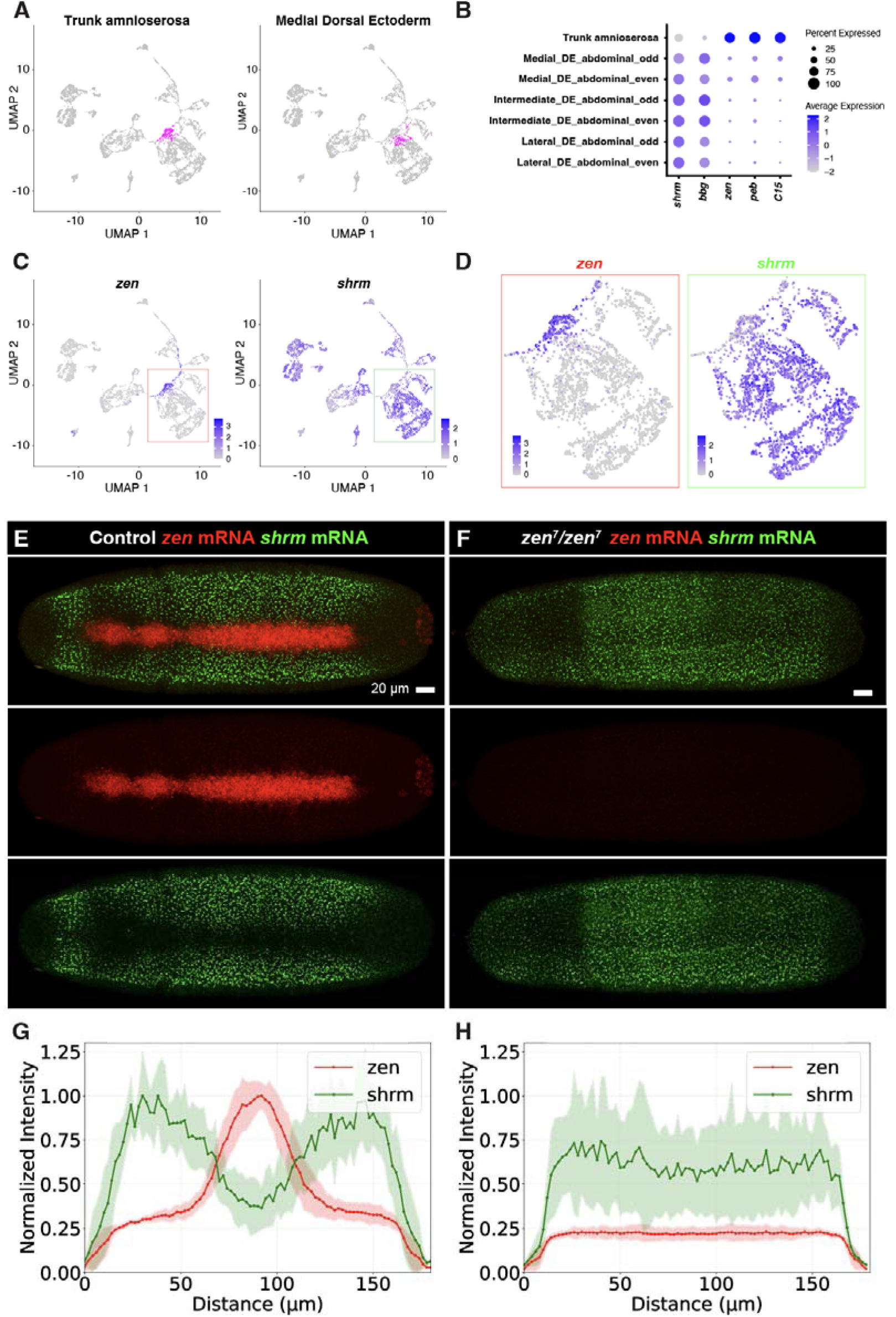
Zen transcriptionally silences *shrm* in the AS. (A-D) Re-analysis of the scRNA-seq data from Sakaguchi et al. 2023^40^. (A) Cells annotated as trunk_amnioserosa (left) or medial dorsal ectoderm (medial_DE_abdominal_even and medial_DE_abdominal_odd; right) are highlighted in magenta on the UMAPs. (B) Dot plot showing the expression patterns of representative genes that are specifically expressed or repressed in trunk amnioserosa cells. (C) Expression of *zen* (left) and *shrm* (right) on the UMAPs. (D) Magnified views of the boxed regions in (C). (E and F) Dorsal view of a control (E, n=3) or a *zen* mutant (F, n=3) embryo showing HCR of *zen* (red) and *shrm* (green) transcripts. Scale bars, 20 µm. (G and H) Spatial intensity profiles of *zen* and *shrm* transcripts measured at 50% EL in the dorsal region of control (G, n=3) or *zen* mutant (H, n=3) embryos. See also Data S2 and S3.

To verify the scRNA-seq prediction that Zen represses *shrm*, we performed hybridization chain reaction (HCR) to visualize *shrm* mRNA. We found that *shrm* mRNA is present in the ectodermal cells outside the AS primordium in a pattern largely complementary to that of *zen* mRNA (Fig. 5E, 5G). In *zen* mutants, *shrm* mRNA is de-repressed in the dorsal midline cells, resulting in a uniform profile of distribution across the entire dorsal half of the embryo (Fig. 5F, 5H). These data confirm that Zen transcriptionally represses *shrm* in the AS.

We next examined whether Zen-dependent repression of *shrm* accounts for low junctional myosin in the AS cells. We performed a double RNAi knockdown of *zen* and *shrm*. While in the dorsal cells E-Cad levels are comparable between *zen shrm* double and *zen* single RNAi embryos (see Limitations of the study), the levels of junctional myosin are significantly lower in *zen shrm* double RNAi than in *zen* RNAi alone (Fig. 6A-C). These data support the hypothesis that Zen-dependent transcriptional repression of *shrm* ensures low levels of junctional myosin in the AS cells. They also suggest the possibility that tissue rupture seen in the *zen^-^* embryos results from Shrm-dependent elevation of junctional myosin contractility. In support of this, we observed partial suppression of tissue rupture phenotype in *zen shrm* double knockdown (3/6 embryos show rescue of tissue rupture, see also Limitations of the study). We therefore predict that ectopic expression of Shrm in the AS cells could be sufficient to increase junctional myosin, and thus cell-intrinsic stresses, thereby causing tissue rupture.

**Figure 6.**
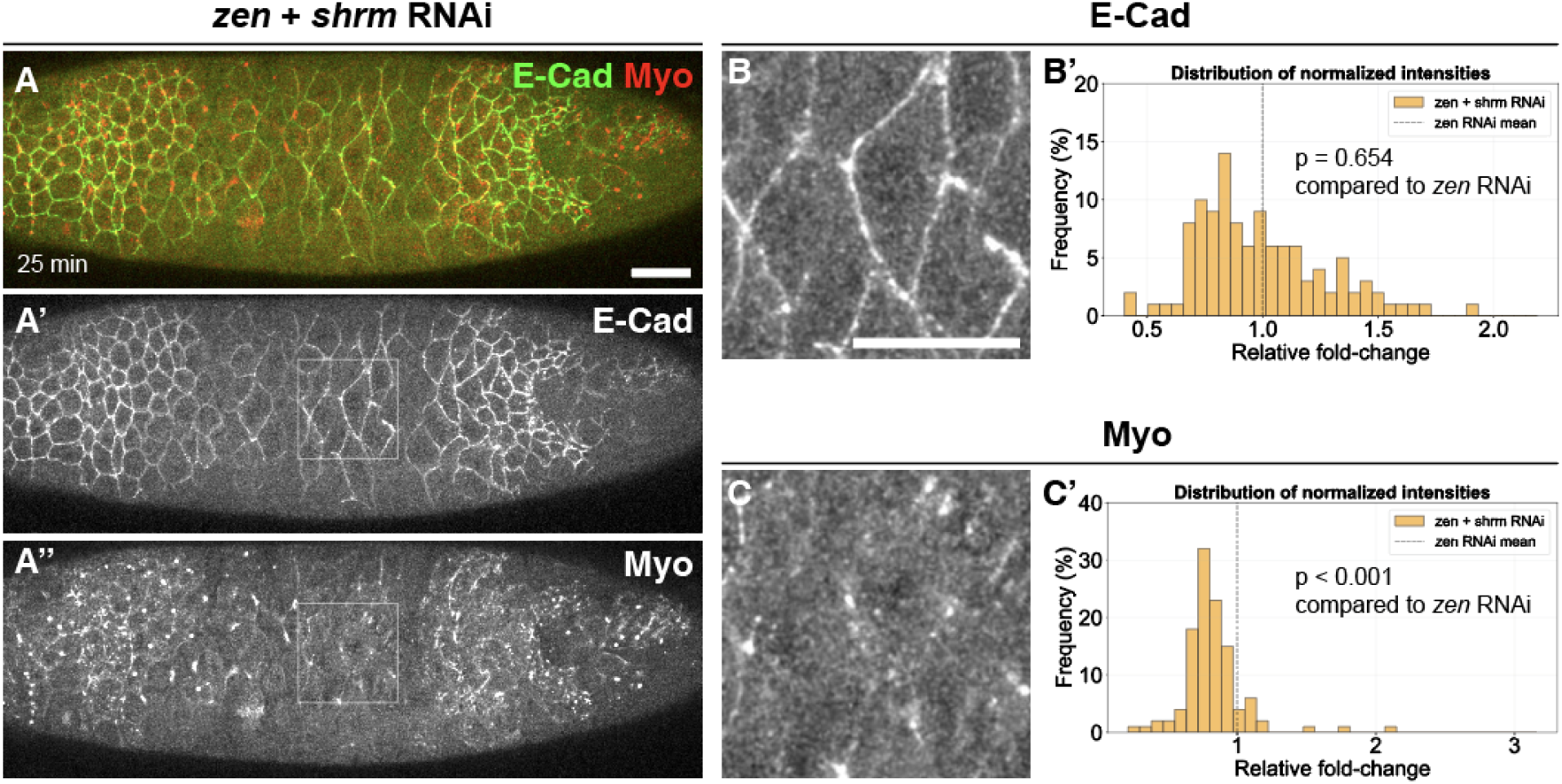
Up-regulation of junctional myosin in *zen* RNAi requires *shrm*. (A-A’’) Dorsal view of a *zen shrm* double RNAi (n=3) embryo expressing E-Cad-3xGFP (Green in A; A’) and MyoII-3xmKate2 (Red in A; A’’), 25 min after the onset of AS apical elongation. White boxes outline a representative region of the AS enlarged in B and C. (B and C) Enlarged view of regions outlined in white in A’ and A’’, highlighting a decrease of junctional myosin in *zen shrm* double RNAi (C), as compared to *zen* RNAi alone (Figure 4F). Scale bars, 20 µm. (B’ and C’) Histogram showing the distribution of E-Cad and myosin junctional intensities, 25 min after the onset of AS apical elongation in *zen shrm* double RNAi (B’ and C’, n=96 junctions). Mann-Whitney U test; p = 0.654 (B’) and p < 0.001 (C’).

### MitoTRAP Gal4: an optogenetic Gal4 system that delivers on-demand, spatially targeted ectopic expression of target genes

To test this prediction, we sought to elevate *shrm* expression levels in the AS cells. Because an AS-specific Gal4 driver is currently unavailable, we set out to develop a light-inducible Gal4 system that can allow for on-demand, spatially targeted expression of target genes. We reasoned that expressing the full complement of Gal4 (Gal4DBD+VP16AD) as a single protein, while tethering it to the organelles that are compartmentized even before the completion of cellularization, could allow us to ensure the spatial restriction of Gal4 nuclear translocation following photo-activation, even if we start photoactivation during cellularization (see Methods for further discussion of the design rationales). To achieve this, we created the mitoTRAP Gal4 system (Fig. S3A). In brief, full length Gal4-VP16 was fused with LANS4, which contains the photo-sensitive LOV2 domain that cages a nuclear localization signal (nls), and the resultant Gal4-VP16-LANS4 was co-expressed with the mitochondrion-anchored Zdark tag (dNTOM20-Zdk2) of the LOVTRAP system^44–46^. In the dark, we expect dNTOM20-Zdk2 to sequester Gal4-VP16-LANS4 to the mitochondria. Following blue light illumination, conformational change of LOV2 is expected to allow Gal4-VP16-LANS4 to become dissociated from Zdk2, while the uncaging of the nls in LANS4 could facilitate the nuclear translocation of Gal4-VP16-LANS4 to drive the transcription of the downstream target gene.

To assess the efficacy of the mitoTRAP Gal4 system, we expressed it maternally and assayed its performance in the embryo by quantifying the zygotic expression of UASp-EGFP-nls. With the flies and the embryos both kept from blue light prior to imaging, we first varied the copy number of dNTOM20-Zdk2 to assay the mitochondrion sequestration efficacy. We imaged EGFP-nls in a collection of embryos at varying stages from the onset of cellularization, which marks the onset of zygotic transcriptional activation, to the successive timepoints during gastrulation, staged according to PMG position. In embryos containing Gal4-VP16-LANS4 alone, we observed that the levels of EGFP-nls increase with developmental time, indicating that in the absence of dNTOM20-Zdk2, Gal4-VP16-LANS4 drives the expression and accumulation of EGFP-nls efficiently (Fig. S3B). In contrast, in embryos containing either a single copy or two copies of dNTOM20-Zdk2, the expression of EGFP-nls is consistently low from cellularization through to late-gastrulation (Fig. S3B), indicating that dNTOM20-Zdk2 functions as an effective trap that prevents Gal4-VP16 from accessing the nuclei.

We next quantified the kinetics and spatial restriction of EGFP-nls expression following localized photoactivation. Quantitation reveals that EGFP-nls intensity increases with time specifically in nuclei associated with the photoactivation ROI, but not nuclei elsewhere (Fig. S3C, C’). These data show that the mitoTRAP Gal4 system can be used to effectively activate target gene expression with high degrees of spatial specificity and with activation kinetics compatible with the timescale of AS morphogenesis.

### Ectopic expression of Shrm in the AS is sufficient to cause tissue rupture

We then proceeded to use the mitoTRAP Gal4 system to ectopically activate *shrm* in the AS cells. Quantitative analysis reveals that spatially restricted activation of *shrm* expression in the AS cells leads to increased junctional myosin (Fig. S4A, A’’, B, B’’, E, F), suggesting that ectopic Shrm is sufficient to raise junctional contractility and cell-intrinsic stresses. Interestingly, we also observed a decrease in the levels of E-Cad (Fig. S4A, A’, B, B’,C, D), which may account for low E-Cad levels observed in the *zen^-^* embryos. We next asked whether mitoTRAP Gal4 driven UAS-Shrm expression leads to tissue rupture. To infer the site of UAS-Shrm ectopic expression, we included UASp-EGFP-nls as a reporter to visualize cells in which the mitoTRAP Gal4 system has become active following photo-activation. We observed tissue rupture consistently (3/4 embryos), and these ruptures are similar to those observed in the *zen^-^* embryos, with the sites of rupture spatially coinciding with EGFP-nls expression (Fig. 7A, Movie S6). Thus, ectopic, local expression of *shrm* in the AS is sufficient to cause tissue rupture in a manner similar to the removal of Zen, thus establishing a causal link between Shrm expression and tissue rupture. In contrast, ectopic expression of Shrm via the mitoTRAP Gal4 system in the AS cells is unable to cause tissue rupture in *bnt* mutants (3/3 embryos; Fig. 7B, 7C, Movie S7), substantiating that tissue rupture depends on external stresses. Taken together, these results provide a direct support for our model that Zen transcriptionally represses *shrm* to lower junctional myosin and cell-intrinsic stresses, thereby conferring mechanical compliance to the AS cells in such a way that external stresses can be effectively released to prevent tissue rupture.

**Figure 7.**
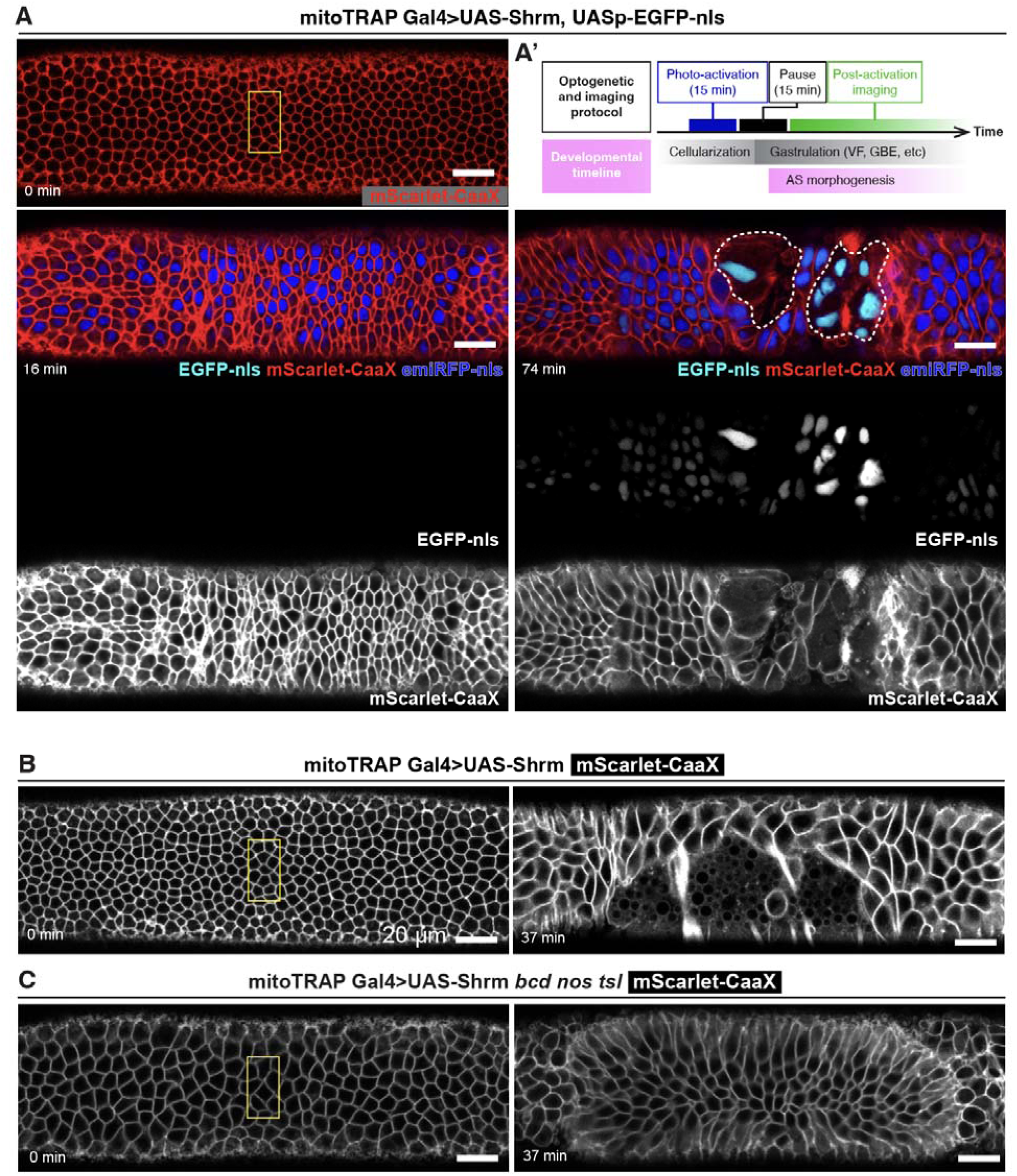
Ectopic expression of Shrm in the AS causes tissue rupture in the presence of external stresses. (A and A’) Dorsal view of an embryo expressing the mitoTRAP Gal4 system employed to drive ectopic expression of UAS-Shrm with co-expression of UASp-EGFP-nls as the reporter to visualize the region of expression. Yellow box, region of photo-activation; white dotted outline, rupture areas. Time labels are elapsed time from the end of photo-activation. The experiment protocol for the photo-activation (see also Methods) is depicted in (A’). 3xmScarlet-CaaX (red and gray) and emiRFP-nls (blue and gray) are used to visualize cell membranes and nucleus. EGFP-nls (cyan and gray) visualizes the cells expressing the UAS transgenes. (B and C) Dorsal view of an embryo expressing the mitoTRAP Gal4 system for ectopic expression of Shrm in the wild-type (B) and *bnt* mutant (C) backgrounds. Yellow box, region of photo-activation. Time labels are elapsed time from the end of photo-activation. 3xmScarlet-CaaX is used to visualize cell membranes. Scale bars, 20 µm. Related to Figures S3, S4, Movie S6 and S7.

## Discussions

In this study, we reveal a previously unknown mechanical function for the *Drosophila* extraembryonic tissue, the AS, during gastrulation. Specifically, AS squamous morphogenesis releases tensile stresses yielded by VF formation and GBE, two morphogenetic events that produce large-scale contractile forces that temporally overlap with AS morphogenesis. Previous studies have shown that VF formation and GBE exert traction forces on the neighboring tissues to drive large-scale tissue flows^3,27,28^. Pulling forces generated by VF formation have additionally been shown to be transmitted across tens of micrometers to bias the orientation of mitotic spindles in the head ectoderm^47^. Although AS surface expansion accounts for less than 3% of the lost surface area, contrasting with cell division that accounts for ∼65%^12^, loss of the AS leads to large-scale tissue ruptures, while gastrulation morphogenesis appears largely normal in *cdc25/string* mutants, despite the reduced cell number^11,31^. Thus, we suggest that AS surface expansion functions as the major dissipator of tensile stresses, while in the absence of it, contractile forces generated by these large-scale tissue flows are highly detrimental to gastrulation.

AS morphogenesis performs its function as a stress dissipator by being mechanically compliant. Such pliability requires transcriptional repression of *shrm*. Shrm is an evolutionary conserved scaffolding protein known to promote actomyosin contractility via recruitment of Rho-kinase to the actin-rich membrane cortex or junctions^41–43^. It either instructs myosin-dependent apical constriction, or is required for the establishment or maintenance of myosin planar polarity, or both, in a wide range of naturally occurring morphogenetic processes, including neural tube closure, lens placode invagination, *Drosophila* GBE, as well as optogenetically induced deformation of cells in an organoid model^41,42,48–52^. Our experiments of ectopically expressing Shrm in the AS cells via the mitoTRAP Gal4 system causally link the increased junctional myosin to tissue rupture. In these experiments, as well as in the *zen^-^* embryo, where we observed tissue rupture, the AS cells are initially stretched by the external pulling forces, showing an increase of strain. By the time when the tissue tears began to form, the AS has all but halted its apical surface elongation, suggesting that rupture does not result from a continuous increase of externally induced strain, but is due to increased intrinsic stresses resulting from the rise of Shrm expression that drives the increase of junctional myosin. Thus, as briefly alluded to in Results, these observations serve as an *in vivo* confirmation for the recent *in vitro* data that reveal a regime of epithelial rupture in a substrate-free culture of Madin–Darby canine kidney (MDCK) cells. Specifically, administering Calyculin to a MDCK monolayer that is pre-stretched to a constant strain results in an increase of internal stresses initially and rupture of the tissue eventually as the stresses reach the tensile strength of the tissue^36^. Notably, both this *in vitro* and our *in vivo* systems form tissue tears internal to the tissue, and not near the outer boundary, underscoring the possibility that the underlying mechanics is similar.

In addition to increased intrinsic stresses, we note, however, that weakened cell-cell adhesion might also contribute to tissue rupture given that junctional E-Cad is lower in the *zen^-^* embryo or with ectopic expression of Shrm than in the wild-type. Shrm is not known to be directly involved in the control of junctional E-Cad localization in *Drosophila* embryo, nor is it in the mammalian epithelia. As such, the apparent inhibitory effect of Shrm on E-Cad might be indirect. In the neighboring ectodermal epithelium, Shrm is known to maintain planar polarized contractility by promoting junctional enrichment of Rho-kinase during GBE^42^. Given that Rho-kinase phosphorylates and facilitates the delocalization of Bazooka/Par-3, a scaffolding protein that recruits E-Cad to adherens junctions, it is conceivable that Shrm indirectly represses junctional E-Cad by elevating Rho-kinase junctional recruitment, which in turn reduces Bazooka/Par-3 levels^53^. A dorsal-to-ventral gradient of E-Cad localization was recently reported and attributed to EGFR signaling, although the mechanism is not known^54^. On the contrary, junctional myosin has been shown to form a reversed, ventral-to-dorsal gradient that is thought to be related to Dpp/BMP signaling^3,28,55,56^, which is upstream of Zen. It would thus be of interest to see whether the spatial patterning of Shrm expression contributes to the Rho-kinase/myosin gradient and secondarily to the E-Cad gradients along the dorso-ventral axis.

That the AS morphogenesis functions as a stress dissipator during gastrulation adds on to other known functions of the AS in orchestrating late-stage morphogenetic events, such as germband retraction ^57–59^ and dorsal closure^60–62^. In thinking of how these morphogenetic functions arose during evolution, it is of critical importance to note that the AS is a derived state in Diptera that originated in the stem group of the late-branching schizophora, which includes *Drosophila*. In contrast, most dipterans outside schizophora, and in fact most other insects, form two extraembryonic tissues – the serosa and the amnion – that represents the ancestral state^63–67^. The serosa tissue undergoes an epiboly-like expansion towards the ventral side to ultimately wrap around the entire embryo, forming a bilayered tissue topology between the enveloping serosa, and the amniotic-embryonic epithelium^65,68,69^. Given this tissue topology, it would be of interest to see how external tensile stresses generated by mesoderm invagination and ectoderm convergent extension are dissipated, and in particular, whether *shrm* is transcriptionally repressed in the serosa, or in the amnion, or both. Of note, the serosa in the non-schizophoran flies and other insects undergoes a late-stage rupture at late stages after it has enveloped the embryo as part of the process of extraembryonic withdrawal to allow the fully developed larva to break through these outer extraembryonic tissue layers to hatch^63^. Tissue rupture has been shown to be an essential component in a growing number of morphogenetic processes^70^, and in particular, increased contractility is implicated in a number of processes featuring tissue rupture^71–73^. It would thus be of particular interest to see whether the serosa rupture is in fact driven by increased contractility, and more specifically, by a temporal-spatially controlled upregulation of Shrm expression that elevates cell-intrinsic stresses to induce rupture.

In sum, that AS squamous morphogenesis functions as a dissipator of tensile stresses supports and complements our recent work on the CF, where we showed that CF out-of-plane deformation pre-empts the accumulation of compressive stresses resulting from the collision of head and trunk ectodermal tissues, thereby preventing tissue buckling^11^. Programs such as these mitigate deleterious effects of mechanical conflicts, which could arise during development, especially when multiple morphogenetic events take place concurrently, and when the events themselves are insufficiently compartmentalized in space. We propose that the evolution of the dissipator function of AS squamous morphogenesis might fall within the conceptual framework that proposed recently, namely physical constraints or mechanical conflicts might prime species to evolve dedicated, specialized programs of mechanical stress management^11^. A deeper understanding of how the stress dissipator function of the amnioserosa arose via comparative and phylogenetic explorations of extraembryonic tissue evolution in species that branched just before the evolutionary onset of this function, will be of critical importance in testing and enriching this conceptual framework.

## Limitations of the study

We showed that both cell-intrinsic compliance and external tensile stress function as mechanical components contributing to AS squamous morphogenesis. These, together with previous work that emphasizes on the remodeling of the microtubule network as a cell-intrinsic process driving AS elongation^20^, suggest that AS squamous morphogenesis involves both active and passive deformations. While this conclusion is in line with a wide range of diverse work that reveals the contribution of the external stresses, or the intrinsic remodeling of cytoskeleton, or both^74,75,17,76–78^, in orchestrating squamous morphogenesis, neither have we exhaustively identified all sources of external stresses, nor has the list of candidate Zen targets (Data S2 and S3) been sufficiently tested to provide a more comprehensive picture of how AS active deformation is controlled cell autonomously. Thus, there remains much to be done to understand the forces and mechanisms involved in AS squamous morphogenesis.

The possibility that other unknown Zen targets are involved in conferring mechanical compliance to the AS cells might be one explanation for why the *zen shrm* double RNAi embryos show only a partial rescue (50%) of tissue rupture. Another technical caveat of these experiments is that the loss of *shrm* could potentially decrease stresses yielded by GBE^42^. Furthermore, there remain possibilities that one or more uncharacterized Zen target genes directly involved in elevating cell-cell adhesion (Data S2 and S3), in addition to the indirect influence that we speculate for Shrm. With these complications involved with the *zen shrm* double RNAi experiments, we think that the ectopic expression of Shrm via the mitoTRAP Gal4 system remains the more definitive evidence that causally links Shrm to tissue rupture in the *zen^-^* embryo.

---

**Figure S1.**
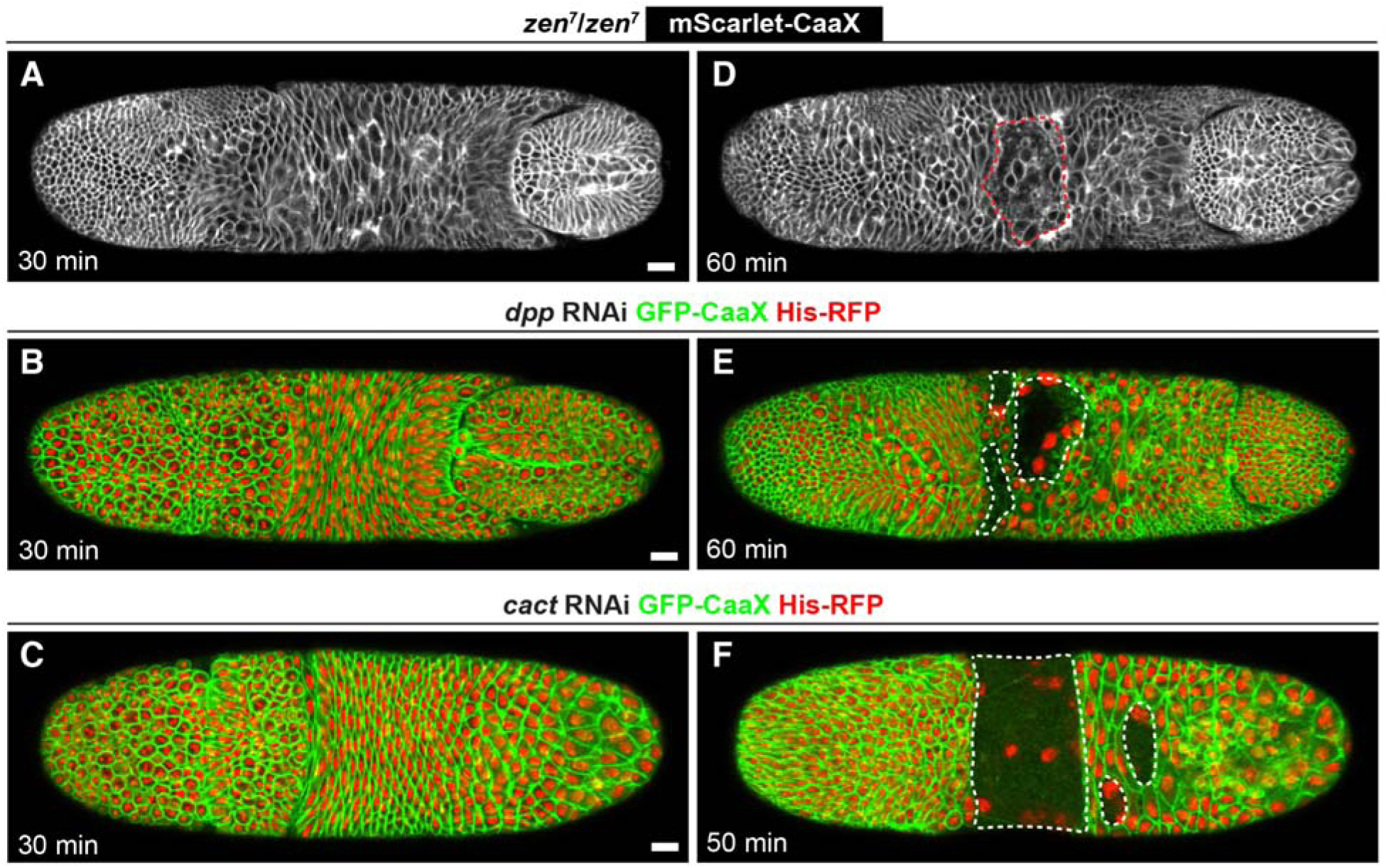
Loss of *zen* or its upstream regulators results in similar phenotypes of partial AS apical elongation and eventual rupture of the AS tissue. (A-F) Dorsal view of a *zen* mutant (n=3) embryo expressing 3xmScarlet-CaaX, or a *dpp* RNAi (n=3) or a *cact* RNAi (n=3) embryo expressing GFP-CaaX and His-RFP, at 30 min (A-C) and 50 or 60 min (D-F) after the onset of AS apical elongation. Red or white dotted outlines, rupture areas. Scale bars, 20 µm. Related to Figures 2 and 3.

**Figure S2.**
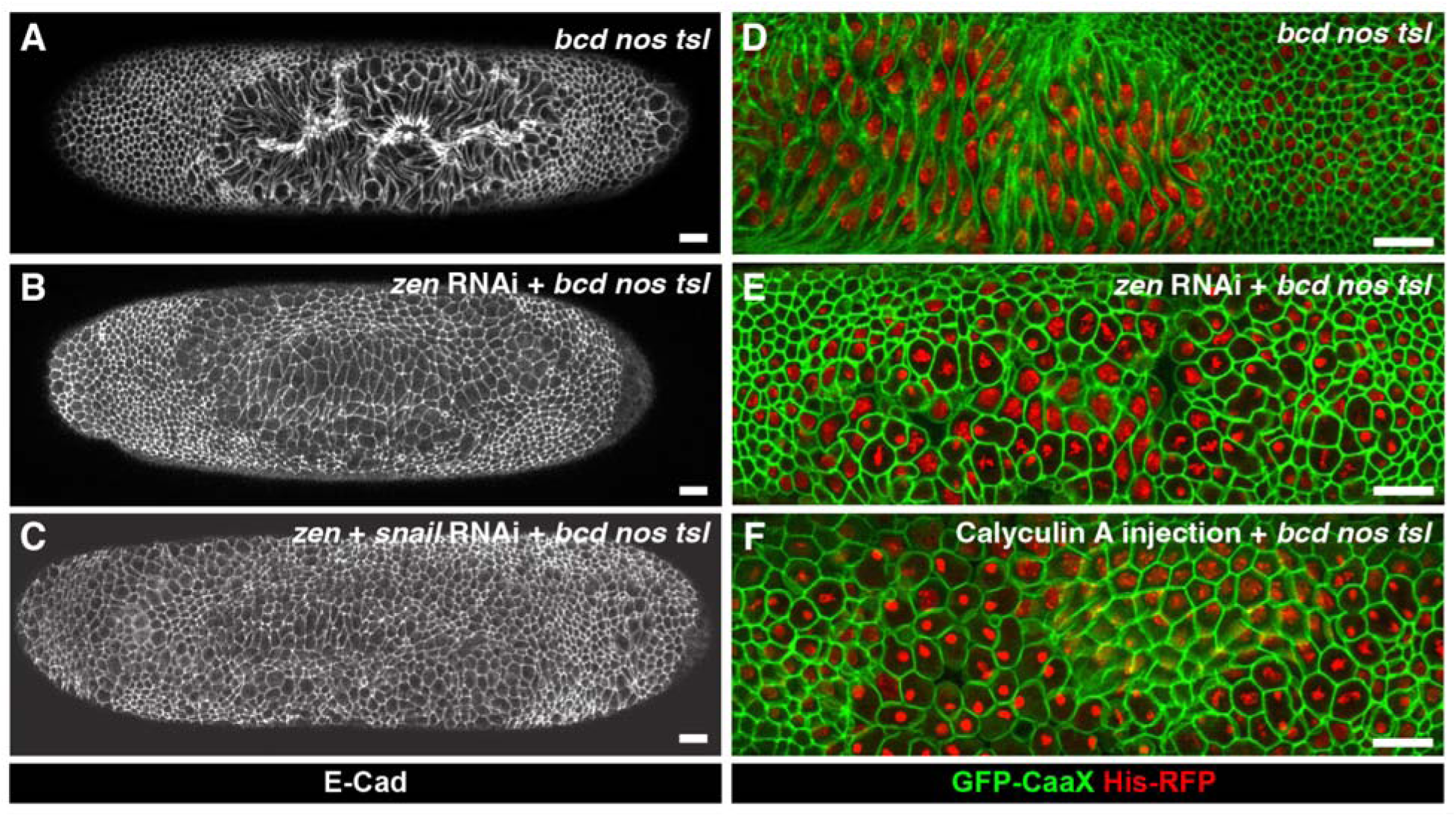
AS apical elongation requires external stresses and low intrinsic contractility. (A-C) Dorsal view of a control *bnt* mutant (A, n=3), or a *bnt* mutant with *zen* RNAi (B, n=3), or a *bnt* embryo with *zen snail* double RNAi (C, n=3) expressing E-Cad-3xGFP, showing a progressive loss of AS apical elongation as the AS fate (*zen* RNAi), and the AS fate plus VF formation (*zen snail* double RNAi) are lost, in conjunction with the loss of GBE (*bnt* mutant). (D-F) Dorsal view of a control *bnt* mutant (D, n=3), or a *bnt* mutant with *zen* RNAi (E, n=3), or a *bnt* embryo injected with Caluculin A (F, n=3), expressing GFP-CaaX and His-RFP, showing that an increase in actomyosin contractility (Calyculin injection) can inhibit AS apical elongation to an extent comparable to *zen* RNAi. Scale bars, 20 µm. Related to Figures 2 and 4.

**Figure S3.**
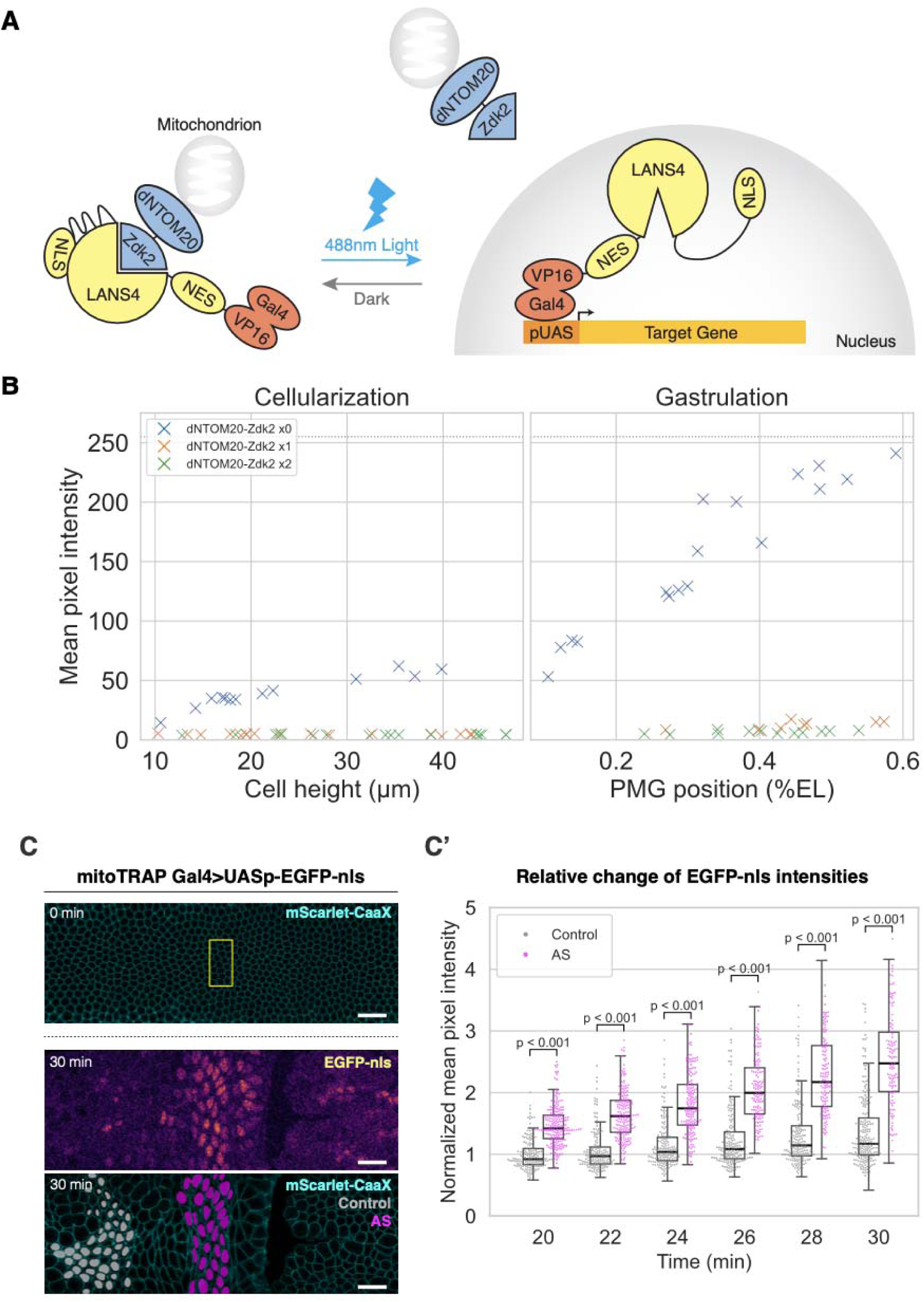
Characterization of the optogenetic Gal4 system: mitoTRAP Gal4. (A) Schematic illustration of the mechanistic principle of mitoTRAP Gal4. In dark, Zdk2 binds to LANS4, thereby trapping Gal4-VP16 at the mitochondrion. With 488nm light illumination, conformational change of LANS4 leads to its unbinding from Zdk2 and the uncaging of nls, allowing Gal4-VP16 to be translocated to the nucleus, and thereby activating the transcription of target gene downstream of UAS. (B) The efficacy of mitochondrion trapping via dNTOM20-Zdk2. Gal4-VP16-LANS4 was co-expressed with 0 copy (blue, n=30 embryos), 1 copy (orange, n=26 embryos), or 2 copies (green, n=29 embryos) of dNTOM20-Zdk2, and its transcriptional activity in the dark was essayed using UASp-EGFP-nls. Mean pixel intensities of nuclear EGFP plotted as a function of development stages in cellularizing (measured by the cell height, left subplot) and gastrulating (measured by PMG position, right subplot) embryos. (C and C’) Representative images of the embryo of mitoTRAP Gal4 driving UAS-EGFP-nls (C) at the end of photo-activation (0 min; cyan, 3xmScarlet-CaaX; yellow box, the ROI of photo-activation), and 30 min after photo-activation, showing the EGFL-nls channel alone (top; inferno look-up-table), or the segmented and binarized nuclear masks with the 3xmScartletCaaX channel (bottom; grey, nuclear masks for the control region in the head; magenta, nuclear mask for the AS region; cyan, 3xmScarlet-CaaX). See Methods for further details for how these two regions were selected. Swarm and box plots (C’) of nuclear GFP intensities showing the increase of nuclear GFP intensities in the AS relative to the control regions (AS: n=109∼194 nuclei; control: n=1158∼209 nuclei; from 3 embryos) as a function of time after photo-activation. Nuclear GFP intensities are normalized by the mean value of the control region at 20 min after activation. The box plot shows the central horizontal line as the median, the box as interquartile range, and the whiskers as the minimum and maximum data points excluding the outliners. Mann-Whitney U test; p < 0.001. Related to Figure 7.

**Figure S4.**
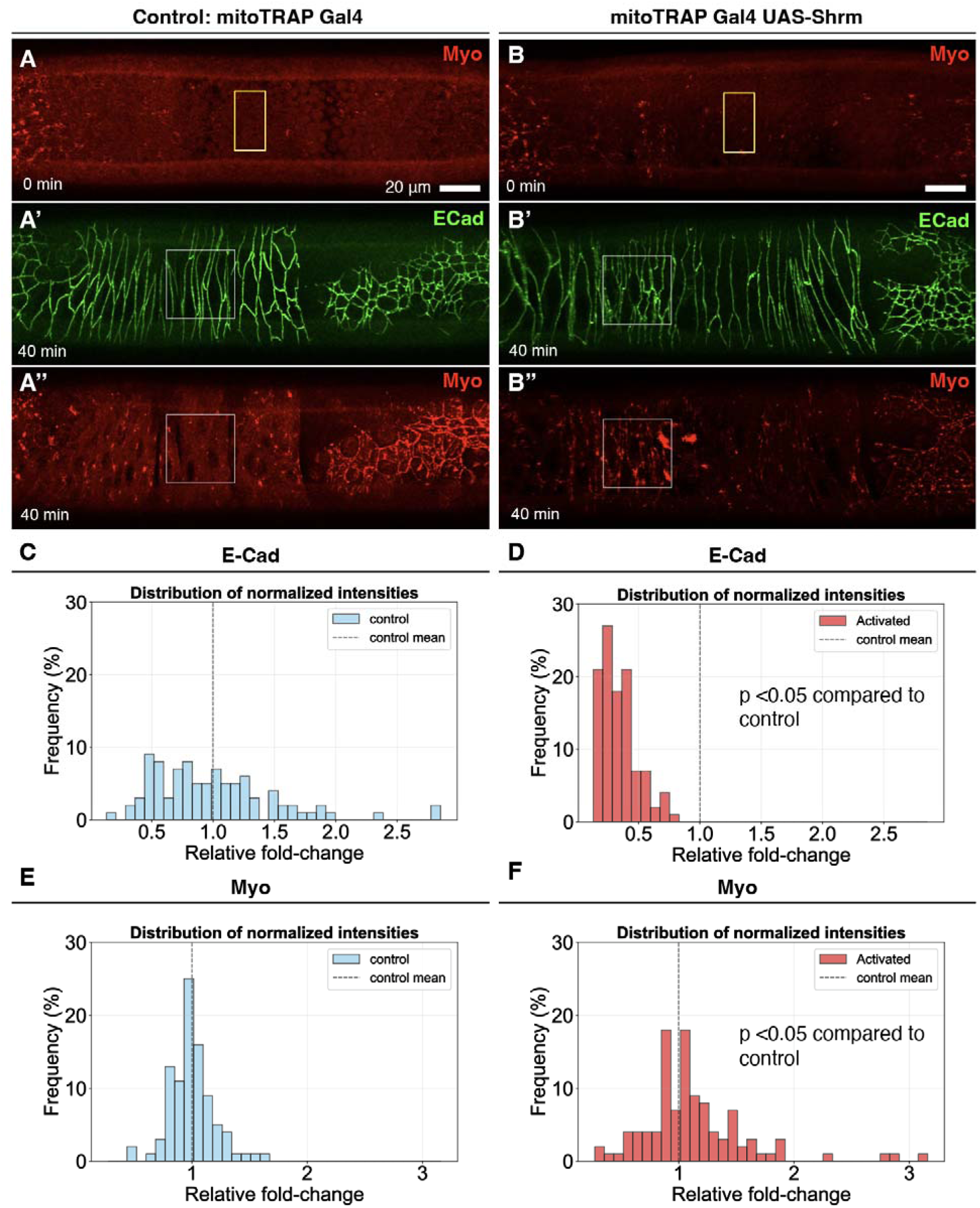
Ectopic expression of *shrm* elevates junctional myosin. (A and B) Dorsal view of a control embryo (A-A’’, n=3) or an embryo expressing the mitoTRAP Gal4 system for ectopic expression of UAS-Shrm (B-B’’, n=3). Each embryo was subject to an identical photo-activation regiment (see Methods) in the region outlined by the yellow box (A and B), followed by E-Cad-3xGFP (A’ and B’) and MyoII-3xmKate2 (A’’ and B’’) imaging, shown here at 40 min after photo-activation when the levels of junctional myosin appear up-regulated (white boxes) in the Shrm-overexpressed embryo, as compared to control. Scale bars, 20 µm. (C-F) Histograms showing the distribution of E-Cad and myosin junctional intensities, measured at 40 min after photo-activation for individual junctions from the regions outlined in the white boxes in (A’, A’’, B’, B’’), in control (C and E, n=80 junctions) and Shrm overexpression (D and F, n=98 junctions). Compared to control, mean junctional E-Cad intensity in Shrm overexpression is significantly lower (D), while mean junctional myosin intensity (E) is significantly higher. Mann-Whitney U test; p < 0.05. Related to Figure 7.

**Figure S5.**
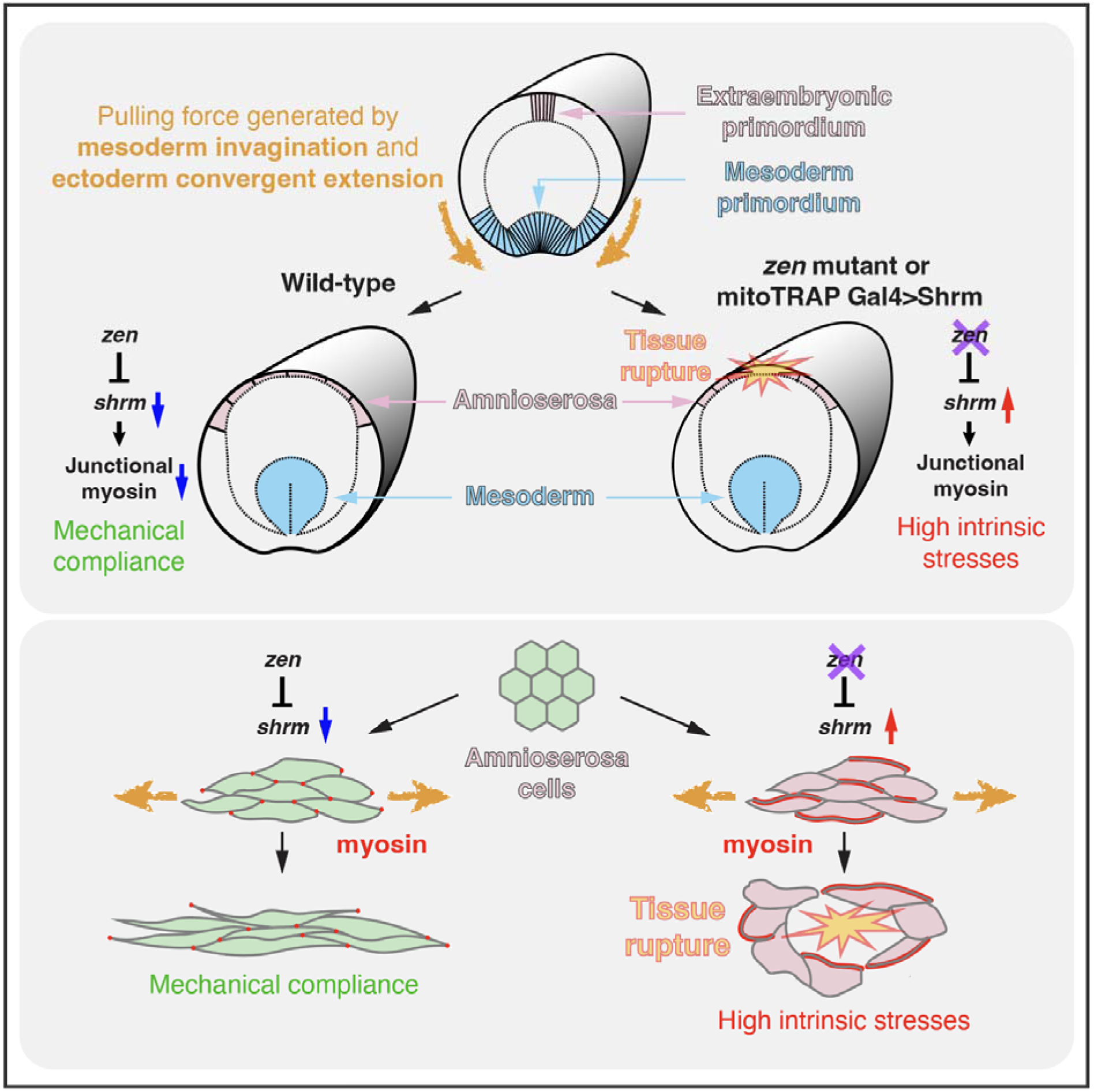
Schematic model for the release of tensile stresses via cell-intrinsic mechanical compliance in the AS. Pulling forces generated by mesoderm invagination (VF formation) and ectoderm convergent extension (GBE) exert on the extraembryonic primordium cells at the dorsal midline, yielding an increase of tensile stresses. In the wild-type embryo, the Zen-dependent transcription silencing *shrm* allows AS squamous morphogenesis to sufficiently release such external stresses. Conversely, in *zen* mutants, or in the embryos with ectopic expression of Shrm in the AS, insufficient release of external tensile stresses in conjunction with an increase in intrinsic stresses resulting from Shrm-dependent recruitment of junctional myosin leads to tissue rupture.

## Movie legends

**Movie S1**

Multi-view morphodynamics of the timing of AS morphogenesis in relationship to VF formation and GBE. A cross-sectional view (A0) and a lateral surface projection view (A) of the same embryo, and a dorsal surface projection view of separate embryo (B) temporally aligned based on PMG position, visualized with GFP-CaaX. Time labels are elapsed time from the onset of gastrulation defined by the onset of VF formation (arrowhead). Related to Figure 1.

**Movie S2**

Dorsal view of a control, *zen* RNAi, *bnt* mutant, *bnt* mutant with *zen* RNAi, *zen* and *snail* double RNAi or *stg* mutant with *zen* RNAi embryo visualized with GFP-Caax and His-RFP or 3xmScarlet-Caax showing AS morphogenesis and the eventual tissue rupture that occurs in selected genotypes. Time labels indicate elapsed time from the onset of AS apical elongation. Related to Figures 2 and 3.

**Movie S3**

Dorsal view of a *bnt* mutant, *bnt* mutant with *zen* RNAi, or *zen* and *snail* double RNAi embryo visualized with E-Cad-3xGFP showing a progressive decrease of AS apical elongation. Related to Figure 2.

**Movie S4**

Dorsal view of the AS region in a *zen* RNAi embryo expressing E-Cad-3xGFP and 3xmScarlet-CaaX showing tissue rupture in single z-slice views at an apical (left) and subapical (right) planes. Time labels indicate elapsed time from the start of imaging. Related to Figure 3.

**Movie S5**

Dorsal view of a control, *zen* RNAi, or *zen* and *shrm* double RNAi embryo showing the differences in junctional E-Cad-3xGFP and MyoII-3xmKate2 intensities during AS morphogenesis. Time labels indicate elapsed time from the onset of AS apical elongation. Related to Figure 4.

**Movie S6**

Dorsal view of an embryo expressing 3xmScarlet-CaaX and emiRFP670-nls for the localized tissue rupture phenotype resulting from UAS-Shrm expression following spatially restricted photo-activation of mitoTRAP Gal4. UASp-EGFP-nls is used to visualize the region of mitoTRAP photoactivation. Time labels indicate elapsed time from the onset of AS apical elongation. Related to Figure 7.

**Movie S7**

Dorsal view of a control, or a *bnt* mutant embryo (top) expressing 3xmScarlet-CaaX for the tissue rupture phenotype resulting from UAS-Shrm expression following spatially restricted photo-activation of mitoTRAP Gal4. In contrast to the control, tissue rupture is suppressed in the *bnt* mutant embryo (bottom). Time labels indicate elapsed time from the onset of AS apical elongation. Related to Figure 7.

## Supplementary datasets

**Data S1:** Time of initial rupture relative to the onset of dorsal cell elongation in all genotypes observed.

**Data S2:** The list of AS high genes and the associated statistics.

**Data S3:** The list of AS low genes and the associated statistics.

## Methods

### *Drosophila* genetics and transgenic lines

*Drosophila* lines used for live imaging were GFP-CaaX^79^, His-RFP (FBtp0056035), E-Cad-3xGFP^80^, MyoII-3xmKate2^80^, 3xmScarlet-CaaX^79^ and emiRFP670-nls (this work). *Drosophila* mutant alleles used were *zen*^7^ (FlyBase ID: FBal0018861), *stg^7M53^* (FlyBase ID: FBal0016176), and the *bnt* triple mutant that contains *bicoid^E1^* (FlyBase ID: FBal0001080) *nanos^BN^* (FlyBase ID: FBal0044220) *tsl^3^* (FlyBase ID: FBal0017197). Identification of *zen* or *stg* homozygous mutant embryos in live imaging experiments was based on the absence of a balancer-linked reporter construct, hb0.7-Venus-NLS, inserted on the TM3 balancer^21^. The emiRFP670-nls was generated by cloning the emiRFP670^81^ (addgene#136556) appended at the C-terminus with the nls from P{UAS-Stinger} (FBtp0018187) into pBabr-mat-tub-(insert)-3’UTR, a ΨC31 site-directed transformation vector, that contains the mat-tub promoter (gift from D. St. Johnston, Gurdon Institute, UK) and the spaghetti-squash 3′ UTR.

For *cactus* and *dpp* RNAi, females of Transgenic RNAi Project stocks, *P{TRiP.HMS00084}^attP2^* (*cact*) and *P{TRiP.JF02455}^attP2^* (*dpp*), were crossed to males of *matαTub-Gal4VP16^67C^; matαTub-Gal4VP16*^15^ double driver line that also contains imaging markers described above. The resultant F1 flies were used to set up egg deposition cages that were kept at 18°C for collection of embryos used in the experiments.

### dsRNA synthesis

For *zen, snail* and *shrm* RNAi experiments, double stranded RNA (dsRNA) was synthesized on templates that contain the T7 promoter sequence (5’-TAATACGACTCACTATAGGGAGA-3’) at each end using a MEGAscript T7 kit (Ambion); templates were amplified from 0–4 h embryonic cDNA using specific primers (5’-CTGCCACATCTAGCACAGGA-3; 5’-AGACGGTTGCTTAGCTCCAA-3) for *zen*, (5’-CGGAACCGAAACGTGACTAT-3; 5’-GCGGTAGTTTTTGGCATGAT-3) for *snail* and (5’-TTGAACAACACTGATCCCGA-3; 5’-TTCGAACTTTTGCCGTAACC-3) for *shrm*.

### Injections

For dsRNA injections, 0-1 hr old (up to stage 2) embryos were collected, dechorionated with bleach and mounted on an agar pad. The mounted embryos were then picked up using a coverslip painted with glue (prepared by immersing bits of scotch tape in heptane), desiccated for 10-14 min using Drierite (W. A. Hammond Drierite Co.) and covered with a mixture of Halocarbon oil 700 and 27 (Sigma-Aldrich) with a ratio of 3:1. Needles for injection were prepared from micro-capillaries (Drummond Microcaps, O.D. 0.97 mm, I.D. 0.7 mm) pulled with a Sutter P-97/IVF and beveled with a Narishige pipette beveller (EG-44). Injections were performed on a Zeiss Axio Observer D1 inverted microscope using a Narishige manipulator (MO-202U) and microinjector (IM300). A volume of ∼100 pL solution was injected into the embryos with a concentration of 1-1.5 μg/μl for dsRNA. After injection, embryos were kept at 25°C injection in a moist chamber until mid to late-cellularization, followed by live imaging.

For Calyculin A injections, a volume of ∼100 pL solution with a concentration of 0.1 mg/ml Calyculin A (Sigma-Aldrich, 208851) was injected into embryos just before the onset of gastrulation, followed by live imaging.

### Live imaging

Embryo-scale imaging for relating AS morphodynamics to other gastrulation morphogenetic events was performed on a two-photon scanning microscopy with a 25X water immersion objective (N.A.=1.05) on an upright Olympus FVMPE-4GDRS system (InSight DeepSee pulsed IR Dual-Line laser, Spectra Physics) with an excitation wavelength at 930 nm for EGFP-CaaX in two imaging angles: lateral and dorsal. Embryos were collected, dechorionated, mounted on glass-bottom dishes, and immersed in water for imaging. Live imaging of the dorsal surface was performed on a Leica SP8 confocal microscope using a 20X (N.A.=0.75) or 63x (N.A.=1.40) oil immersion objective. Z stacks were typically acquired with a ∼20 µm depth, image size of 246.2 x83.8 µm and a z step size of 0.9 µm with a 1 min time interval for a dorsal view of the embryo. High speed imaging was done using the Yokogawa CSU-W1 SoRa Confocal Scanner Unit. Z stacks were typically acquired with a ∼5.6 µm depth, image size of 96.6 x96.6 µm and a z step size of 0.5 µm with a 20 s time interval for a dorsal view of the embryo.

### Reanalysis of single-cell RNA-seq data

Identification of AS-specific genes using previously constructed gastrula-stage scRNA-seq data^40^ was performed as follows. First, the Seurat object data was obtained from https://data.mendeley.com/datasets/k8g638cmxv/1 and loaded using Seurat (version 4.4.0). Differentially expressed genes between trunk_amnioserosa and medial_DE_abdominal (DE=dorsal ectoderm) cells were then identified using the FindMarkers function (FindMarkers(ident.1 = “trunk_amnioserosa”, ident.2 = c(“medial_DE_abdominal_even”, “medial_DE_abdominal_odd”), test.use = “MAST”)) with adjusted p-values based on bonferroni correction and a threshold of 0.01. Plots of the scRNA-seq data were also generated using Seurat.

### MitoTRAP Gal4: design principle, construction and execution

We initially experimented with the recently published shine-Gal4 system^82^, but were not able to achieve spatially restricted expression (data not shown). The shine-Gal4 system uses the photo-switchable Magnet, with the DNA binding domain (DBD) and the transactivating domain (AD) of Gal4 expressed as two independent peptides that separately fused to each of the two Magnet monomers that dimerize following blue light illumination. We suspect that the lack of spatial restriction may be due to the diffusion of the photoactivated monomers across cell units that have not been fully compartmentalized prior to the completion of cellularization.

We thus employed a double-gating design emulating a recent report^83^, where the dark-state mitochondria trapping based on the LOVTRAP (LOV2 Trap and Release of Protein) system^46^ was combined with a light-inducible nuclear translocation approach^45,84^. We called this optogenetic Gal4 system “mitoTRAP Gal4”. Specifically, Gal4-VP16 is fused to LANS4 (Light Activatable Nuclear Shuttle)^45,84^, an engineered photo-sensitive LOV2 domain that cages a nuclear localization signal (NLS), and co-expressed with a mitochondria-anchored Zdark tag (dNTOM20-Zdk2) from the LOVTRAP system for mitochondria anchorage^46^. Prior to photoactivation, Gal4-VP16-LANS4 is expected to be anchored at the mitochondria via binding of its LOV2 domain to dNTOM20-Zdk2 with high affinity. Following blue light illumination, conformational change of LOV2 allows Gal4-VP16-LANS4 to be dissociated from Zdk2, while the NLS in LANS4 become uncaged to facilitate its nuclear translocation (Figure S3A).

To generate MitoTRAP Gal4 fly lines, DNA fragments containing Gal4-VP16, mScarlet, NES, and LANS4, dNTOM20 and Zdk2 were custom synthesized (Eurofins, Japan), and inserted into pBabr-mat-tub-(insert)-3’UTR to generate pBabr-Gal4-VP16-mScarlet-NES-LANS4 and pBabr-dNTOM20-Zdk2. These were then integrated on Chromosome II at ZH-51C and VK00037, respectively, or on Chromosome III at ZH-86Fb and attP2, respectively, by phiC31 site-directed integration (WellGenetics, Taiwan).

To execute optogenetic activation of UAS target genes, female flies that contain the mitoTRAP Gal4 system and the imaging markers were crossed to males containing UASp-EGFP-nls (P{UAS-Stinger}3, FBti0127871), or UAS-Shrm (P{UAS-Shrm.B}3, FBti0167128), or both. To prevent unwanted photoactivation, fly crosses, cages for egg deposition, and embryos prior to processing were kept in the dark. Prior to imaging, embryos were processed, staged and mounted in a dark room with the desk lamp and the brightfield light source on the stereo microscope both covered by a light red filter (#027, Lee filters, UK). Imaging was performed on a Leica SP8 confocal microscope using a 63x oil immersion objective (N.A.=1.40). Photo-activation of the mitoTRAP Gal4 system started during early cellularization when the cell height was ∼20 μm (Figure S3C) or ∼13 μm (Figures S4 and 7), and was achieved using the FRAP mode with a 488-nm Argon laser set at 3% laser intensity. For each iteration of photo-activation, an ROI of 16x32 µm positioned at the mid-dorsal area covering the amnioserosa region was illuminated for 3 times with a duration of ∼8 s, followed by a Z stack imaging with a 561 nm laser for 3xmScarlet-CaaX (or MyoII-3xmKate2, Figure S4, or with a 633 nm laser for emiRFP670-nls, Figures S3C and 7A) to monitor the progression of development for a duration of ∼59 s. The above iteration was repeated for 15 times, which amounted to a total duration of ∼15 min. Following the photo-activation, the embryo was imaged for 3xmScarlet-CaaX (or MyoII-3xmKate2, Figure S4, or emiRFP670-nls, Figures S3C and 7A) for a duration of ∼20 min (Figure S3C) or ∼15 min (Figures S4 and 7), during which the exposure to 488 nm light was paused. Following the pause, post-activation imaging ensued, with the use of 488 nm laser to image the EGFP-nls reporter for data shown in Figure S3C and Figure 7A, or E-Cad-3xGFP for data shown in Figure S4, alongside the imaging of 3xmScarlet-CaaX (or MyoII-3xmKate2, Figure S4, or emiRFP670-nls, Figures S3C and 7A). Note for data shown in Figures 7B and 7C, only the 561 nm laser was used for imaging of 3xmScarlet-CaaX, but not the 488 nm laser.

### Hybridization Chain Reaction

For *zen* and *shrm* mRNA detection, Hybridization Chain Reaction (HCR) was used. Embryos were fixed by heat-formaldehyde method. Probes were synthesized by Molecular Instruments (Los Angeles, CA. USA). Probe set size was 20. Transcripts were detected fluorescently as described (dx.doi.org/10.17504/protocols.io.bunznvf6).

### Image processing and quantification

Images were processed, assembled into figures and converted into videos using FIJI or python. Quantitative data were analyzed and processed by custom-made Python scripts using Numpy, Pandas, and SciPy libraries. Plots were generated with Python scripts using Matplotlib and Seaborn graphic libraries. The sections below provide a brief description for each of the image processing and analysis procedures.

#### Surface projection

For *en face* views, the FIJI plugin Local Z Projector^85^ or a custom-made Python implementation of the projection method was used to project a surface of interest from a 3D stack onto a 2D surface, taking into account the curvature of the embryo. The reference plane that represents the contour of the embryo surface was derived from a Gaussian-blurred image of the original 3D stack image with a σ value of 2∼4, followed by binarization with a customized threshold, which results in a smooth height map for z projection

#### Time annotation

The majority of the time-lapse images were temporally aligned and annotated based on the increase of apical surface area and the aspect ratio in the AS cells, which marks the onset of AS elongation. In Figure 1, the onset of gastrulation was defined on the basis that the initiation timings of VF and PMG are concurrent. To align datasets based on VF onset, for each dataset 2∼4 manually tracked VF cleft depths from embryo surface (d_VF_) were fitted linearly to estimate the actual time for when d_VF_=0. To align datasets based on PMG onset, the anterior edge of PMG was manually tracked to measure d_PMG_, the distance between PMG to the posterior pole, and measured distances for the first 15 min were fitted linearly to estimate the actual time for when d_PMG_=0. The averaged time difference between d_VF_=0 and d_PMG_=0 was then used to align the averaged dynamics of PMG positions to that of the VF cleft depths, which then served as the timescale for measurements of AS apical area and aspect ratio. AS apical areas and aspect ratios were measured based on segmentation of 3xmScarlet-CaaX.

#### Measurement of AS apical surface elongation and elongation orientation

Segmentation of E-Cad-3xGFP was performed using the MorphoLibJ plugin in Fiji to obtain the apical surface of individual AS cells. To estimate the extent of cell elongation and elongation orientation, geodesic diameter and Feret angle were measured, respectively, using the MorphoLibJ plugin in Fiji for the segmented apical surfaces.

#### Measurement of junctional E-Cad and myosin intensity

Segmentation of E-Cad-3xGFP was performed using the MorphoLibJ plugin in Fiji to obtain a skeletonized junctional network. Vertices where 3 or more junctions meet were dilated and subtracted from the junctional network to allow isolation of individual junctional masks that separate two neighboring cells. The total intensities of E-Cad-3xGFP or MyoII-3xmKate2 were measured within each junctional mask and divided by the number of pixels in each mask to obtain mean E-Cad or myosin intensities for each junction.

#### Plotting of *zen* and *shrm* intensity profile

Maximum projection of dorsal view of the embryos stained with HCR *in situ* hybridization was used for the measurement. The image was positioned with the embryonic A-P axis parallel to the X axis. Fluorescent intensities were measured in a rectangular ROI with a dimension of (X,Y)=(60,180) µm with the X dimension centered at the 50% EL, while Y dimension centered on the dorsal midline. Averaged pixel intensities along the X dimension were re-scaled to the min and max of the control mean and plotted as a function of distance along the Y dimension of the ROI.

#### Measurement of EGFP-nls in mitoTRAP Gal4 experiments

Surface projection was performed as described above using the Python implementation of local Z projection. The emiRFP-nls channel was then binarized with multi-Otsu thresholding implemented in Scikit-image to generate a single mask of all nuclei (Figure S3B) or with Cellpose-SAM^86^ to generate masks of individual nuclei (Figure S3C). For Figure S3B, 2D analysis was performed with a single Z slice that had the largest nuclear mask selected as a representative Z slice for measurement of the mean EGFP-nls pixel intensity for that embryo. For Figure S3C, 3D analysis was performed by generating a 3D nuclear mask: 2D masks of the same nucleus across the Z axis were linked based on their centroid positions. The resultant 3D nuclear mask was then used to compute the mean EGFP-nls intensity for that nucleus. To select the appropriate nuclei for comparison between the AS and control regions, the following procedures were followed: 1) mitotic cells were first removed based on low emiRFP-nls mean intensities due to nuclear envelope breakdown; 2) nuclei located in the region between the CF and the posterior DF were selected as nuclei representing the AS region; 3) nuclei positioned ∼33 µm anterior to the AS region or more were designated as the control nuclei. Note that nuclei posterior to the AS were located on the ventral side of the embryo during cellularization when photo-activation was executed. These nuclei were not considered because they may have been activated by the scattered 488 nm light when the light traversed through the embryo.

### Statistical Analyses

Python scripts using SciPy library were implemented to conduct Mann-Whitney U test for comparing means between two groups. All of the statistical details of experiments, including the number of experiments (n), which represents the number of embryos unless otherwise noted, were indicated in the figure legends.

### Data and Code Availability

Data and codes developed for data analysis are available upon request from the corresponding authors.

## Acknowledgements

We thank the Bloomington and Kyoto *Drosophila* Stock Centers, Yohanns Bellaïche, Eric Wieschaus for sharing reagents; Shigeo Hayashi for support; Nishita Gattani for initial characterization of the mitoTRAP Gal4 system; members of the Wang, Hayashi, Obata and Yoo laboratories for discussions; Girish Deshpande, Bipasha Dey, Juan Manuel Gomez, Tzu-Yi Huang, Steffen Lemke, and Sameer Thukral for critical reading and comments on the manuscript. This work was supported by the core funding at RIKEN BDR; a Japan Society for the Promotion Science (JSPS) Grant-in-Aid for Early-Career Scientists to C.W.K. (20K15810); a JSPS Grant-in-Aid for Transformative Research Areas (A) to Y.-C.W. and T.K. (22H05167); a Human Frontier Scientific Program (HFSP) Young Investigators grant to Y.-C.W. (RGY0082/2015).

## Author Contributions

Y.-C.W. conceived, designed and supervised the study; C.W.K., M.T., and Y.-C.W. generated the reagents, performed the experiments, analyzed the data, wrote software code, and prepared the visualization; S.S. and T.K. analyzed the transcriptomics data; Y.-C.W. wrote the manuscript and all other authors participated in its editing.

## Additional Information

Supplementary Information (Supplementary Figures, Supplementary Movies and Supplementary Datasets) is available for this paper. Reprints and permissions information is available on the journal website. Correspondence and requests for materials should be addressed to Y.-C.W.

